# The origin and evolution of amphibious hearing in pinnipeds

**DOI:** 10.64898/2026.01.14.699601

**Authors:** James P. Rule, Travis Park, Moganavalli Kattan, Camille Grohé, Roxana Taszus, Stephanie M. Palmer, David P. Hocking, Justin W. Adams, Alistair R. Evans, Ian G. Brennan, Tahlia I. Pollock, Daniela Sanfelice, Felix G. Marx, Naoki Kohno, Martin Sabol, Alexander Stoessel, John J. Flynn, Natalie Cooper

## Abstract

Seals (pinnipeds) are the only mammals that can hear in both air and water. How and when they achieved the ability to negotiate such contrasting auditory media remains unknown. Here, we apply 3D shape and phylogenetic comparative analyses to a large dataset of caniform carnivorans (119 species, 217 specimens) to study the emergence of amphibious hearing in pinnipeds despite significant evolutionary constraints. We find support for the cavernous tissue as a functional and evolutionary mechanism for amphibious hearing. This tissue, which fills with blood during diving to equalise air pressure in the ear, enables a shift from in-air to underwater hearing by matching the acoustic impedance of the ear to that of the surrounding water. Early diverging freshwater pinnipeds had impaired hearing underwater. The first marine pinnipeds could hear amphibiously but were limited by a functional tradeoff between hearing abilities and the need to prevent damage from loud underwater sounds. Subsequently, otariids (eared seals) and phocids (true seals) independently acquired middle ear adaptations that expanded their underwater hearing range. This iterative evolution likely facilitated the exploration of novel auditory adaptive zones by crown pinnipeds, resulting in rare acoustic abilities like ultrasonic singing, vocal learning, and keeping rhythm.

## Introduction

Mammalian ears are fundamentally adapted to transmit and amplify sounds in air ^1–3^. Under water, however, their hearing is impaired by a pressure differential across the ear drum that limits sound transmission through the middle ear ^4,5^. Amphibious hearing is challenging in principle, and attendant tradeoffs occur when ancestrally terrestrial lineages become (semi)aquatic ^6,7^. Despite this, extant pinnipeds (true seals, eared seals, and walruses) can hear clearly, localise sound sources, filter useful signals from background noise, and communicate in both environments ^8–13^.

Little is known about how sound reaches the pinniped inner ear underwater ^14–17^. Bone conduction, where sound signals travel through head tissue directly to the cochlea, is often invoked as a potential pathway ^18–20^. However, this mechanism results in a substantial loss of underwater sound signal (30 decibels), is found throughout mammals, and fails to explain both directional hearing and the clarity of perceived sounds ^21^. An alternative hypothesis, that underwater sound may be transmitted via a blood-filled ‘cavernous sinus’ (henceforth referred to as cavernous tissue), has received minimal attention ^17,18,21^. The cavernous tissue is a vascular structure in the ear canal and middle ear of all extant pinnipeds that engorges with blood to equalise air pressure during diving^17,22–25^. Its impedance resembles that of water, which may create a preferential sound pathway ^17,26^. Similar cavernous tissues may facilitate underwater hearing in penguins ^27,28^.

Like its mechanics, the evolutionary origins of amphibious hearing in pinnipeds – and, thus, their complex underwater communication ^29–31^ – remains poorly understood. When did seals first become capable of hearing underwater? And did they, like cetaceans ^6,7^, face a tradeoff with their in-air hearing capabilities ^16,18,32^, at least initially? New fossil descriptions ^33–36^ and recent advances in pinniped systematics ^37–39^ now offer the chance to tackle such questions on a broader scale.

Here, we compile the largest dataset yet assembled of the ears of caniform carnivorans, including living and extinct pinnipeds. We investigate: 1) whether proposed adaptations for amphibious hearing within pinnipeds differ from their nearest terrestrial relatives; 2) when and how amphibious hearing evolved; and 3) how pinnipeds balance the physical constraints of two markedly different auditory environments.

**Figure 1.**
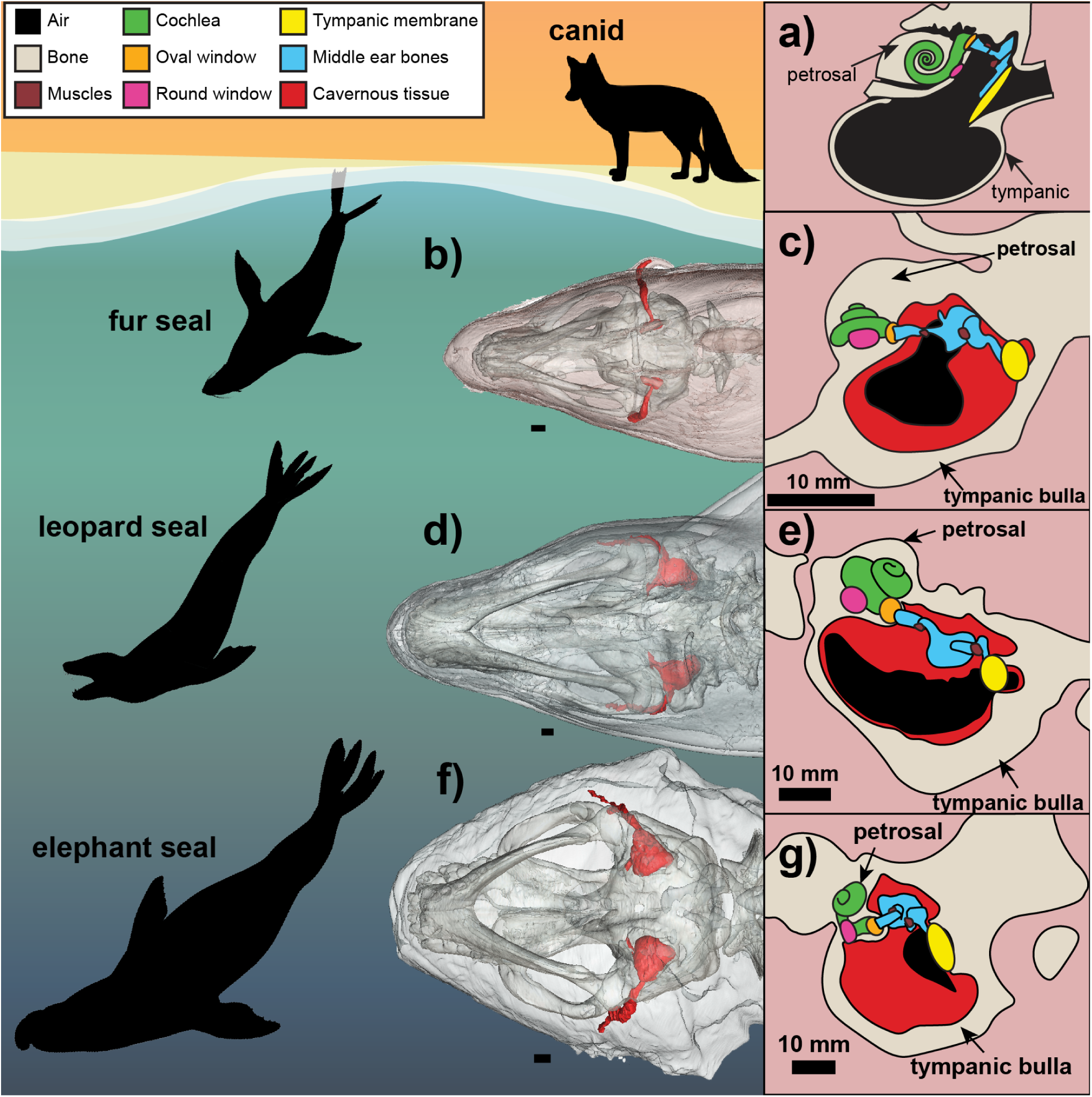
Adaptations of pinniped ears for diving and hearing. The cavernous tissue inflates as pinnipeds dive, equalising air pressure in the ear canal and middle ear cavity. This blood-filled tissue has a similar auditory impedance to water, facilitating sound transmission via the middle ear. Comparisons include idealised lateral sections through the ear (right) and, where present, the cavernous tissue segmented out from computed tomography scans (left) for a) a canid, b) and c) an Australian fur seal (*Arctocephalus pusillus*, TMAG A11092), d) and e) a leopard seal (*Hydrurga leptonyx*, TMAG A11091), f) and g) a southern elephant seal (*Mirounga leonina*, TMAG A11090). Silhouettes: Peter Trusler, except canid (authors). Scale bars 10 mm. Colours represent auditory anatomy.

## Results

We used micro-CT scans of 217 canids, ursids, musteloids, and pinnipeds to compile the largest ever dataset on the anatomy of caniform ears (119 species, including 22 extinct pinnipeds; Table S1) ^14,17,40^. Our eight morphometric measurements — impedance matching ratio, cochlear window ratio, petrosal-basicranium bone contact ratio, angle of the round window from the transverse axis of the petrosal, cochlear turn number, scaled cochlear length, cochlear radii ratio, and cochlear height-width ratio — reflect previously proposed adaptations to amphibious hearing (Figure S1 and Supplemental Information). For each extant species, we also recorded habitat (aquatic vs terrestrial), hearing mode (amphibious vs in-air only), and maximum dive depth (five categories) from the literature (Table S2).

### The cavernous tissue

**Figure 2.**
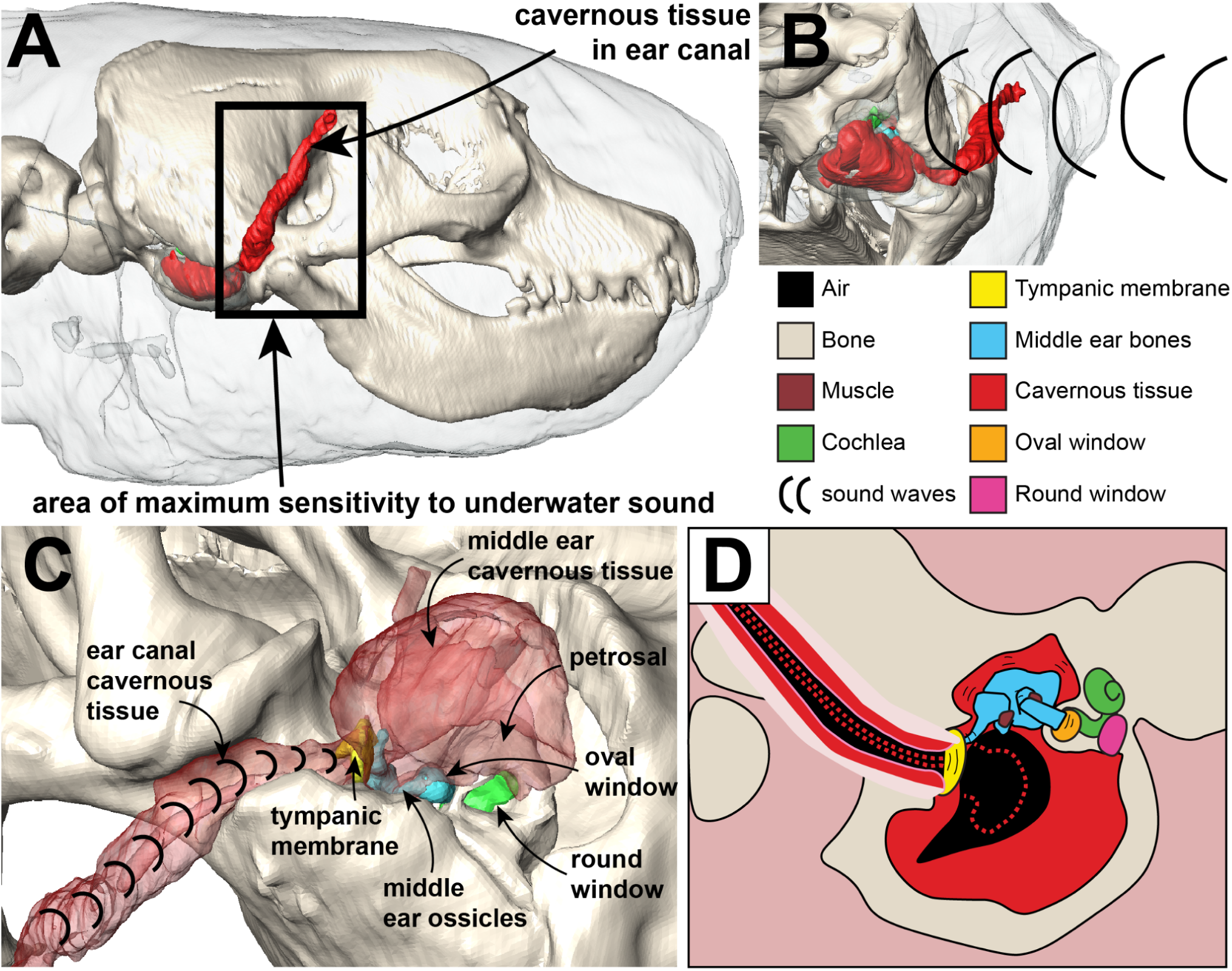
Proposed hearing mechanism for underwater sounds. Illustrated is a southern elephant seal *Mirounga leonina* (TMAG 11090). a) lateral profile of the head (tympanic bulla transparent), showing the distribution of the cavernous tissue in the ear canal, and the area of maximum sensitivity to underwater sounds from Møhl and Ronald ^41^. b) a posterior view of the head, showing a longitudinal soundwave underwater (tympanic bulla transparent). c) a posterolateral view of the head (tympanic bulla removed, cavernous tissue transparent), showing sound waves travelling through the cavernous tissue in the ear canal. d) schematic of the ear, with the hypothesised inflated area of the cavernous tissue indicated by dashed lines. Underwater stimulation of the tympanic membrane (via the cavernous tissue in the ear canal) will result in normal stimulation of the oval window via the middle ear. Figure not to scale.

We segmented out the (deflated) cavernous tissue from soft tissue CT scans of an Australian fur seal (*Arctocephalus pusillus*, TMAG A11092), a leopard seal (*Hydrurga leptonyx*, TMAG A11091), and a Southern elephant seal (*Mirounga leonina*, TMAG A11090). In all three species the cavernous tissue lines both the ear canal and the middle ear cavity, with the auditory ossicles dorsally suspended in sinus tissue (Figure 1). The ear canal and its cavernous tissue run from the auricle behind the eye posteroventrally to the middle ear (Figure 2). The extent of the ear canal lines up with the area of maximum sensitivity to underwater sound for the harp seal from Møhl and Ronald (Figure 2) ^41^. In its deflated state, the tissue abuts the tympanic membrane externally in the fur and leopard seals, and both internally and externally in the elephant seal. When inflated, the tissue could plausibly swell to contact the tympanic membrane internally in all three species. The round window is angled away from the middle ear cavity (postero-ventral in the fur seal, postero-medial in the leopard and elephant seals) and is thus shielded from the cavernous tissue. Previous reports in other pinnipeds ^17,22–24,26,42–44^ indicate that the cavernous tissue is distributed widely throughout Pinnipedia.

One of the main limitations on underwater hearing in mammals is the different acoustic impedance of seawater and air. Acoustic impedance is the opposition of a medium to the transmission of sound, which is determined by differences in the density and speed of sound in those materials. The acoustic impedance of air is 430 kg/m^2^s and seawater is ∼1.56 x 10^6^ kg/m^2^s ^4^, whereas blood is ∼1.61 x 10^6^ kg/m^2^s ^45^. To calculate the reflection fraction of sound travelling between two mediums (with sound travelling in a perpendicular direction) the equation is:

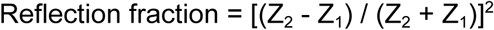

Calculations of impedance demonstrate that 99.9% of sounds are reflected at the seawater-air barrier ^21^, meaning that any air trapped inside the ear canal hinders conduction of sound to the middle ear. When pinnipeds dive, the entrance to their ear canal closes and their cavernous tissue engorges with blood, which has a similar acoustic impedance to seawater. Using the above equation and values, we estimate that <1% of sounds are reflected at the interface between seawater and the cavernous tissue, which makes the latter an efficient pathway for the conduction of underwater sound to the tympanic membrane and, thus, the middle ear. The resulting movement of the auditory ossicles would then stimulate the oval window, as happens during in-air hearing.

Some of the previously proposed osteological correlates of amphibious hearing (e.g. the impedance matching ratio) may indicate hearing through the cavernous tissue.

### Hearing mode, dive depth and habitat are correlated with ear traits

All measurements (Figure S2) except the cochlear radii ratio, window ratio, and petrosal-basicranium contact ratio show strong phylogenetic signal (Pagel’s 𝜆>0.8; Table S3), indicating that traits are similar in closely related species (Supplemental Information). Phylogenetic generalised least squares regressions (PGLS) of log_10_ transformed morphometric data show significant correlations of (i) impedance matching ratio, cochlear window ratio, and petrosal contact ratio with hearing mode; (ii) relative cochlear length, cochlear window ratio, and petrosal-basicranium contact ratio with dive depth; and (iii) the number of cochlear turns, relative cochlear length, and cochlear window ratio with aquatic (pinnipeds, otters, minks, and polar bears) vs terrestrial species (Table 1). These results suggest that aspects of caniform middle ear morphology – including, uniquely, the impedance matching ratio – correlate with hearing mode.

**Table 1.** Phylogenetic generalised least squares (PGLS) regressions of middle and inner ear morphometric data against hearing mode, habitat, and diving category. Traits were log_10_ transformed prior to analyses. 𝜆 = Pagel’s 𝜆. P-values < 0.05 are indicated in bold. Degrees of freedom (hearing mode and habitat) = 1 and 95, degrees of freedom (diving categories) = 4 and 92. Category codings in Table S2. Data plots in Figure S2. Phylogenetic signal tests in Table S3.

| Trait | Hearing mode (in-air, amphibious) |  |  |  | Habitat (Terrestrial, Aquatic) |  |  |  | Diving categories (Terrestrial, Shallow Water, Epipelagic, Mesopelagic, Bathypelagic) |  |  |  |
| --- | --- | --- | --- | --- | --- | --- | --- | --- | --- | --- | --- | --- |
| | F | P | Adjusted R<sup>2</sup> | $\lambda$ | F | P | Adjusted R<sup>2</sup> | $\lambda$ | F | P | Adjusted R<sup>2</sup> | $\lambda$ |
| Cochlear Turns | 3.594 | 0.061 | 0.026 | 0.91 | 4.210 | **0.043** | 0.032 | 0.91 | 1.868 | 0.123 | 0.035 | 0.90 |
| Relative Cochlear Length (mm) | 2.270 | 0.135 | 0.013 | 0.75 | 12.52 | **<0.001** | 0.107 | 0.74 | 5.862 | **<0.001** | 0.168 | 0.75 |
| Cochlear Radii Ratio | 0.008 | 0.931 | -0.010 | 0.41 | 0.050 | 0.823 | -0.010 | 0.42 | 0.277 | 0.892 | -0.031 | 0.38 |
| Cochlear Height-Width Ratio | 1.285 | 0.260 | 0.003 | 0.84 | 0.015 | 0.904 | -0.010 | 0.86 | 1.190 | 0.321 | 0.008 | 0.89 |
| Impedance Matching Ratio | 13.08 | **<0.001** | 0.112 | 0.84 | 2.749 | 0.101 | 0.018 | 0.90 | 1.996 | 0.102 | 0.040 | 0.88 |
| Cochlear Window Ratio | 23.55 | **<0.001** | 0.190 | 0.00 | 0.321 | 0.573 | -0.007 | 0.44 | 5.601 | **<0.001** | 0.161 | 0.00 |
| Petrosal-Basicranium Contact Ratio | 6.278 | **0.014** | 0.052 | 0.37 | 2.277 | 0.135 | 0.013 | 0.51 | 16.75 | **<0.001** | 0.396 | 0.00 |
| Round Window Angle | 3.510 | 0.064 | 0.026 | 0.88 | 0.782 | 0.379 | -0.002 | 0.90 | 1.869 | 0.123 | 0.035 | 0.84 |

### Estimating hearing in extinct pinnipeds

A flexible discriminant analysis (FDA) was trained on the middle and inner ear measurements of the 97 extant caniforms in our data to estimate the hearing mode of 22 extinct pinnipeds. Predictive models using both middle and inner ear data (Figure 3a, Table S4) performed best (FDA resampled accuracy: 98.9%). Crown pinnipeds and stem-pinnipedimorphs (‘enaliarctines’) were estimated to have amphibious hearing, with posterior probabilities >99%. By contrast, the presumed earliest diverging pinnipeds *Potamotherium*† and *Puijila*† were estimated as hearing in air only, with posterior probabilities of 99.61% and 85.64%, respectively. A principal component analysis of the entire dataset (i) distinguishes amphibious from air-only hearing along PC1 (50.5% of sample variation) with contributions from all measurements except cochlear radii ratio (PC2, 13% of variation, is loaded by only cochlear radii ratio) (Figure 3b); and (ii) clusters all extinct pinnipeds except *Potamotherium*† and *Puijila*† with amphibious hearing species. Multivariate analysis of variance of the first seven PC axes (representing 99% of variation) further confirms a significant difference between the two hearing modes (Pillai = 0.89, approximate F = 227, df = 7 and 200, p < 0.001). Equal-rates ancestral states estimation shows that amphibious hearing likely arose in the last common ancestor of crown pinnipeds and ‘enaliarctine’ stem-pinnipeds (Pinnipedimorpha, Figure 3c, Figure S3).

**Figure 3.**
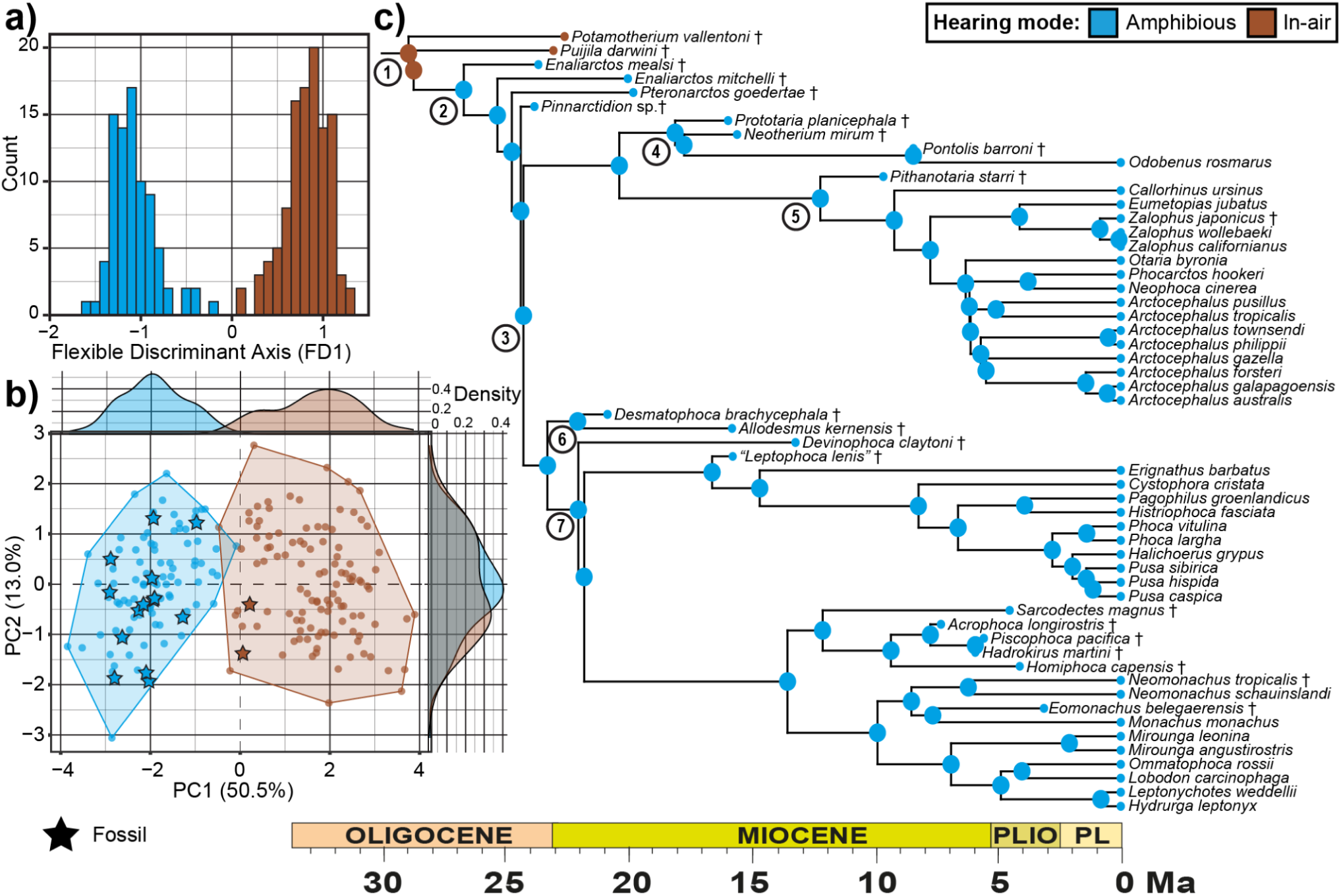
Estimating the origin of amphibious hearing in pan-pinnipeds. **a)** Flexible Discriminant Analysis of middle and inner ear morphometrics distinguishes between air-only and amphibious hearing in caniforms (see Table S4). **b)** Principal Component Analysis of middle and inner ear morphometrics of caniforms, with extinct species represented by stars. **c)** Equal rates ancestral state estimation of hearing mode along the pan-pinniped phylogeny ^38^ (see Figure S3 for full Caniformia ancestral state estimation). Clade names: 1. Pan-Pinnipedia, 2. Pinnipedimorpha, 3. crown Pinnipedia, 4. Odobenidae, 5. Otariidae, 6. Desmatophocidae, 7. Phocidae.

### Evolution of impedance matching in pinnipeds

Our PGLS analyses reveal the impedance matching ratio, i.e. the ratio between the size of the tympanic membrane and the oval window, as a key trait that seems uniquely associated with hearing mode. For mammals that hear only in air, a high impedance matching ratio amplifies sounds to overcome the large impedance difference between air and the cochlear fluid ^1,46^. Underwater, this mechanism is problematic, as the impedance of seawater is more similar to that of cochlear fluid and, thus, could increase sound amplification enough to damage sensory hair cells and nerves of the cochlea ^47–49^.

All crown and ‘enaliarctine’ stem-pinnipeds have comparatively low impedance matching ratios (Table 1, Figure 4), which limit the amplification of underwater sounds to presumably safe levels. This effect is achieved differently between the two largest extant pinniped groups: whereas otariids have a smaller tympanic membrane, phocids instead possess an enlarged oval window (Figure 4a, Supplemental Information). Stem-pinnipeds (*Enaliarctos*†, *Pinnarctidion*†) and desmatophocids show both modifications and, consequently, have a notably lower impedance matching ratio than all other pinnipeds (Figure 4).

All of our morphometric traits show variable rates of evolution suggestive of shifts in the pace of morphological change (Figure 4b, Table S5, Figure S4). Of particular note is a burst in the evolutionary rate of the impedance matching ratio (∼10x the background rate) that coincides with the origin of amphibious hearing along the branch uniting ‘enaliarctines’ and crown pinnipeds (Figure 4b). The angle between the round window and the petrosal also shows multiple, albeit more crownward, rate bursts shortly after amphibious hearing evolved (Figure S4). This may be relevant as a lower angle turns the round window away from the middle ear cavity and, thus, shields it from the cavernous tissue.

**Figure 4.**
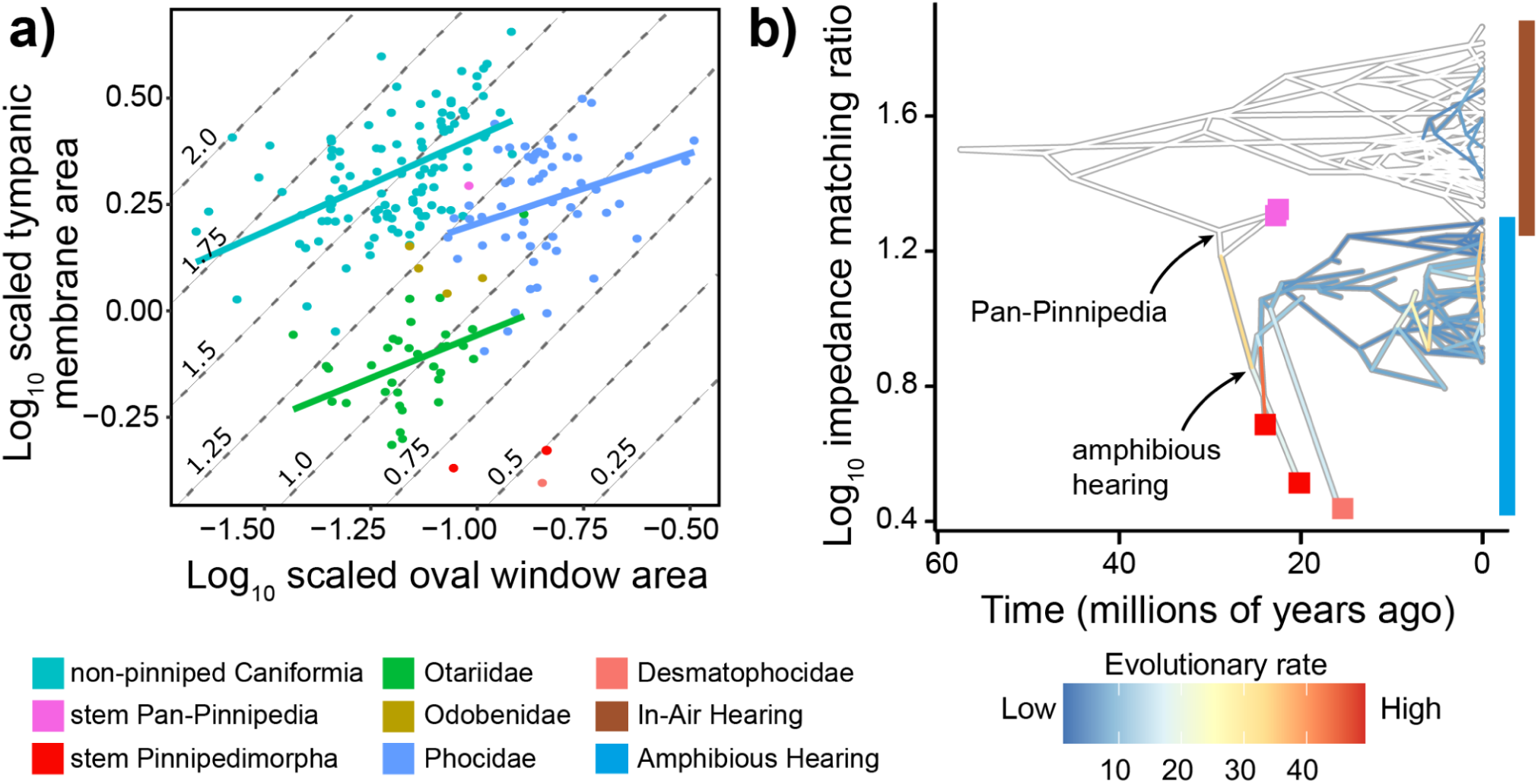
Evolution of the impedance matching ratio in Caniformia. **a)** Scatterplot of log_10_ scaled oval window area against log_10_ scaled tympanic membrane area, with equivalent ratio plotted as contours. Otariids achieve low impedance matching ratios via a small tympanic membrane, whereas phocids have enlarged oval windows relative to the ancestral condition for crown pinnipeds. **b)** Phenogram of variable rates model of log_10_ impedance matching ratio with shifts deviating from the background rate (white branches) mapped on, and relevant fossil tips coloured by clade. See Table S4 for model marginal likelihoods, and Figure S4 for all models.

### Crossing an adaptive valley enabled amphibious hearing

The in-air hearing ranges of pinnipeds are limited compared to their terrestrial relatives (Figure 5), primarily due to a lower High Frequency Hearing Limit (HFHL). In air, otariids have a HFHL that is comparable to ursids and higher than that of phocids and odobenids (Figure 5, Supplemental Information). Extant pinnipeds have far broader underwater hearing ranges than stem-pinnipeds and desmatophocids, with phocids and odobenids notably outperforming otariids (Figure 5).

**Figure 5.**
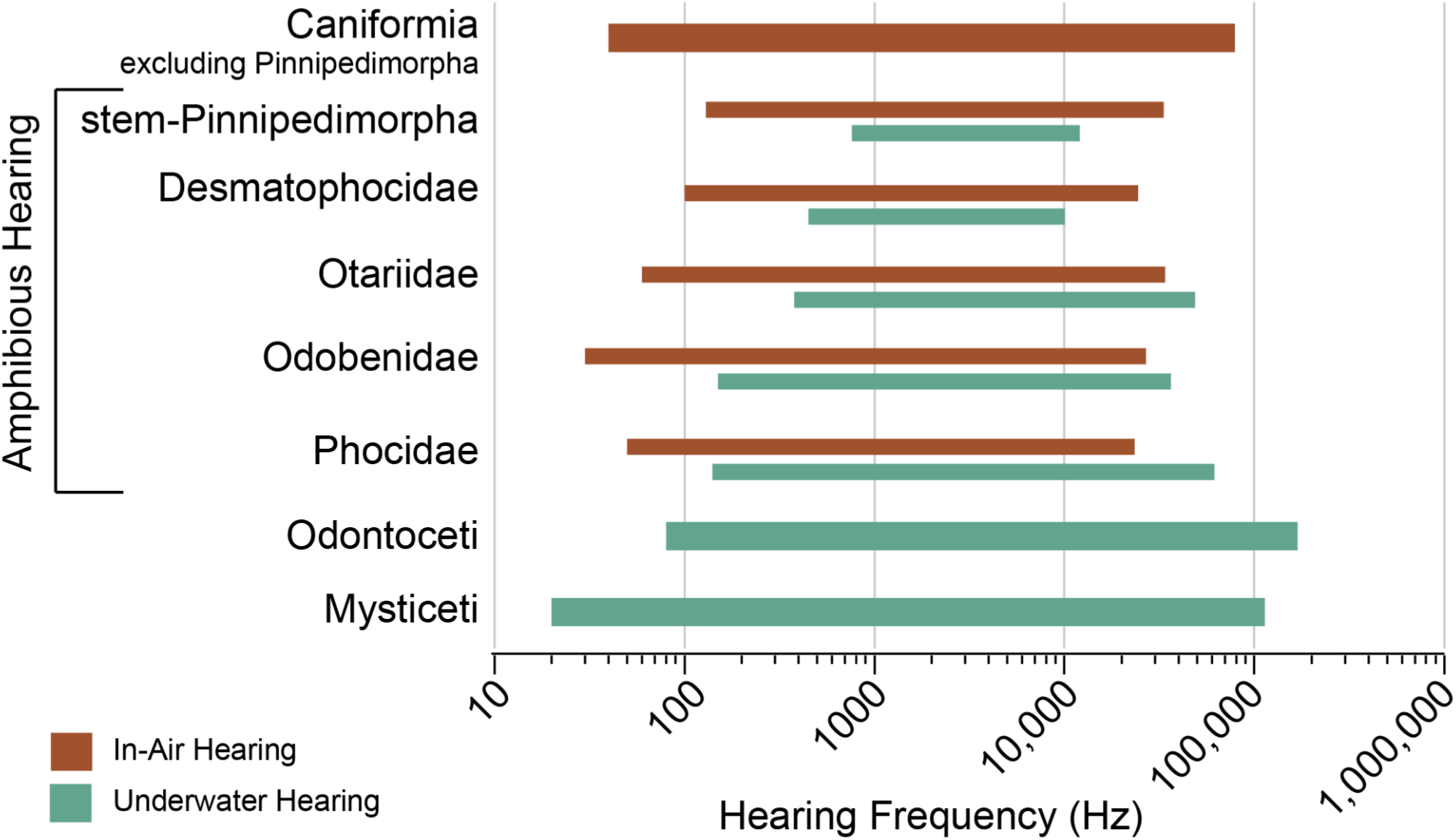
Auditory adaptations for amphibious hearing initially reduced hearing ranges. Relative hearing ranges estimated from inner ear data for Caniformia and extant Cetacea (formula: Supplemental Information and Ref. 50^50^).

We plotted adaptive landscapes synthesising 3D geometric morphometric analyses of cochlear shape and estimated hearing range (to indicate performance) for in-air versus underwater hearing (Supplemental Information) ^40,50^. A PCA of species mean shapes shows limited morphospace occupation (Figure 6, Figure S5), reflecting turn number and relative cochlear length along PC1 (66.7% of variance) and the cochlear height-width ratio along PC2 (21.2% of variance). Eighty theoretical cochlear shapes (represented as landmark sets) were sampled from the morphospace, measured, and hearing ranges calculated from them to construct performance surfaces and adaptive landscapes for hearing mode.

Our in-air hearing adaptive landscape was weighted entirely towards the in-air hearing range (weight = 1), whereas the underwater hearing adaptive landscape had in-air weights of 0.05 and underwater weights of 0.95 to reflect the competing auditory requirements of amphibious hearing species. Pinnipeds occupy lower adaptive values for in-air hearing compared to their terrestrial relatives (Figure 6a), with an adaptive valley coinciding with the transition between air only and amphibious hearing. Several extinct and extant pinnipeds (*Puijilia*†, *Enaliarctos*†, *Neotherium*†, *Pontolis*†, *Odobenus*, *Eumetopias*, and *Sarcodectes*†) occupy this adaptive valley, alongside three ursids, the crab-eating racoon (*Procyon cancrivorus*) and the olingo (*Bassaricyon gabbii*). At the same time most phocids and otariids, the desmatophocids, and the fossils *Prototaria planicephala*† and *Pinnarctidion*† approach an adaptive peak for underwater hearing (Figure 6b). These results suggest that adapting to underwater hearing entailed a notable trade-off in terms of in-air auditory performance.

**Figure 6.**
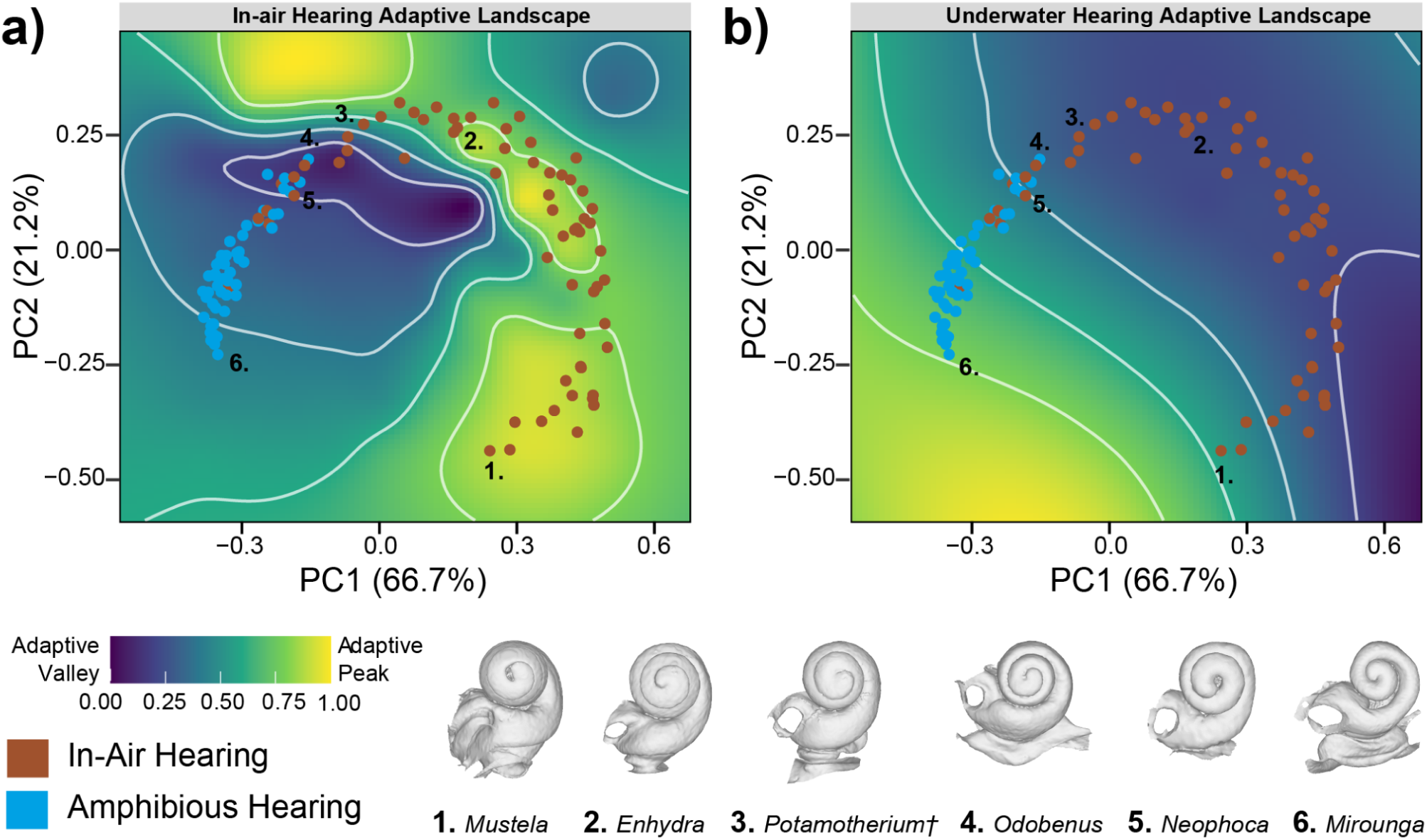
Adaptive landscapes for amphibious hearing. Morphological-auditory adaptive landscapes of a) in-air and b) underwater hearing ranges (in octaves) from 3D geometric morphometric analysis of caniform cochlea (species means). Moving from the terrestrial to the aquatic end of the morphospace (defined by a decrease in coiling and turns of the cochlea), pinnipeds cross an adaptive valley for in-air hearing range, and approach an adaptive peak for underwater hearing range. See Figure S5 for theoretical morphospace and performance surfaces.

## Discussion

Pinnipeds underwent multiple evolutionary transformations during their transition back to the water, including in their locomotor mode ^51,52^, mating systems ^53^, feeding adaptations ^54–59^, life histories ^60^, and sensory morphology ^14,55,61,62^. To date, however, few large-scale studies have included fossils to explore the evolutionary mechanisms underlying sensory transformations ^55,61^.

Our results show that the cavernous tissue of pinnipeds provides a plausible pathway for the conduction of underwater sounds and, thus, amphibious hearing. This idea is consistent with the observation that pinniped heads best receive underwater sounds directly lateral to the cavernous tissue in the ear canal (Figure 2) ^41^. Pinnipeds also have notably lower impedance matching ratios than their non-marine relatives (Figure 4) ^14,17^, which recalls similar middle ear differences between terrestrial and aquatic birds ^28^. Low impedance matching ratios protect the inner ear by reducing the power of underwater sounds and imply that the middle ear is under selective pressure and, therefore, functional during underwater hearing. Together, these findings suggest that pinnipeds hear across their middle ears underwater, directly facilitated by the blood-filled and sound-transmitting cavernous tissue.

The cavernous tissue presumably first evolved to equalise air pressure within the ear during deep diving ^17,22^ and its role in underwater hearing may have been an exaptation. Even so, its appearance likely triggered the evolution of lower impedance matching ratios to limit the amplification of underwater sounds, with several stem-pinnipeds and the desmatophocid *Allodesmus*† showing even lower impedance matching ratios than crown pinnipeds (Figure 4). However, this extreme reduction came at the cost of in-air hearing sensitivity and resulted in relatively narrow underwater hearing ranges (Figure 5), which may ultimately have placed these taxa at a disadvantage.

In modern pinnipeds, vocal communication is critical for fitness signals, competition, territory demarcation, and mother-pup calls, both in-air ^63,64^ and underwater ^29,65,66^. As pinnipeds spread geographically ^38,67,68^ and evolved varied mating systems ^53^, there would have been selective pressure to hear well in both media. A slight secondary increase in the impedance matching ratio, though still below that of terrestrial mammals, would have resulted in correspondingly greater but still manageable amplification of both airborne and underwater sounds, and plausibly would have increased the frequency range of underwater sounds that could be detected. The detailed mechanics of these changes differ between otariids and phocids, which likely explains their disparate in-air and underwater hearing abilities ^18^. The evolution of amphibious hearing in pinnipeds ultimately resulted in auditory adaptations that are rare in mammals, including maintaining rhythm^69,70^, vocal learning ^31,71–73^, and vocal mimicry ^74^.

A better understanding of the function and origins of amphibious hearing can inform how pinnipeds may be impacted by underwater noise pollution, an emerging threat to marine mammals ^47,75–78^. Our findings indicate that amphibious hearing has been important throughout pinniped evolution, and may have originally balanced hearing with minimising harm from loud underwater sounds. Building on this knowledge to understand the hearing abilities of pinnipeds is critical, as harmful sound levels are unknown for most species ^79^.

Transitions to new environments often lead to phenotypic shifts into novel adaptive zones ^80–82^. Land-to-sea transitions among tetrapods are a pertinent example ^83,84^, with marine mammals like cetaceans and sirenians having completely reconfigured their body plan for a marine existence ^84^. Pinnipeds, however, remain partly tied to land, and so must balance adaptations, including to their sensory systems, for operating in both air and water ^37–39^. In our adaptive landscape of in-air hearing (Figure 6a), the transition from air-only to amphibious hearing coincides with an adaptive valley.

Pinnipeds within this adaptive valley include the stem-pinnipeds *Puijilia*† (estimated as being hearing-impaired underwater) and *Enaliarctos*† (estimated as capable of poor underwater hearing), most walruses (except early diverging *Prototaria*†), *Sarcodectes*† (an early diverging monachine phocid), and *Eumetopias* (an extant sea lion with an anomalous hearing threshold in air ^85^).

Unusually, ursids (bears) also occupy and cross this adaptive valley, likely due to their low number of cochlear turns and a potential high frequency range of best hearing ^86^. Other pinnipeds approach, but do not reach, an adaptive peak for underwater hearing (Figure 6b).

Based on our evolutionary modelling, this phenotypic shift towards amphibious hearing occurred during the pinniped adaptation to the marine realm and thus, after their initial transition into freshwater ^37,87^. Extant freshwater carnivorans (otters) occupy adaptive peaks for in-air hearing and adaptive valleys for underwater hearing. Otters neither have nor need cavernous tissue to equalise pressure due to the shallow nature of their dives. As extinct freshwater pinnipeds were likely ecologically analogous to otters, the absence of deep diving would have resulted in limited selection pressure to evolve cavernous tissue, leaving them without a mechanism to hear underwater.

We find that amphibious hearing in pinnipeds has a single origin around 26.7 million years ago, despite the disparate hearing abilities and morphology of otariids and phocids ^15,17,18^. Pinnipeds are unique in being the only extant mammals with amphibious hearing, overcoming specialisations for in-air hearing that evolved over 120 million years ago ^1–3^. This evolutionary innovation may have facilitated the remarkable vocal diversity of pinnipeds ^12,29,64,66^, which – again uniquely among mammals – allows them to communicate both on land and in the ocean.

## Methods

For further details (Methods S1-S4) see the Electronic Supplemental Information.

### Data collection

We compiled a micro-CT dataset (217 specimens; 119 species) for the earbones of extant and extinct pinnipeds and their in-air hearing relatives (Canidae, Ursidae, Musteloidea) from museums worldwide, including previously published datasets ^14,88^. Extinct taxa are denoted with a † symbol in the text. We included inner ear data from 14 cetaceans for comparative purposes (see Supplemental Information). We also micro-CT scanned the wet heads of three pinniped specimens from the Tasmania Museum and Art Gallery collections to segment out the cavernous tissue for

Figure 1. Scans were processed in either Avizo (Thermo Fisher Scientific Inc.) or Dragonfly 3D ^89^, and measurements were taken from the resulting earbone meshes in Meshlab ^90^ and Rhinoceros 8 (Robert McNeel & Associates). We collected the following eight morphological traits for the middle and inner ear ^14,17,40^: (1) the impedance matching ratio (tympanic membrane area / oval window area); (2) the cochlear window ratio (oval window area / round window area); (3) petrosal-basicranium bone contact ratio (full petrosal perimeter / contact between petrosal and basicranium); (4) angle of the round window from the transverse axis of the petrosal; (5) cochlear turn number; (6) scaled cochlear length (cochlear length / geometric mean of cochlea measurements); (7) radii ratio (radius of the outer turn of the cochlea / radius of the inner turn); and (8) cochlear height-width ratio (height of the cochlea / width of the cochlea). See Supplemental Information for details of measurement protocols. Morphological traits were log_10_ transformed, and assessed for normality prior to analyses. For our phylogenetically informed analyses we used a composite phylogenetic tree constructed by grafting non-pinniped trees ^91,92^ onto the pinniped tree of Park et al. ^38^, with the pinniped-musteloid divergence time from Paterson et al. ^37^.

### Analyses

All analyses were run in R version 4.4.2 ^93^ and using the R packages phytools ^94^, mda ^95^, geomorph^96^, and Morphoscape ^82^.

**Phylogenetic statistical tests:** We estimated phylogenetic signal (Blomberg’s K and Pagel’s λ) using the function phylosig for each trait. To test which morphometric traits were important in underwater hearing, we fitted phylogenetic generalised least squares regressions (PGLS) using the R package phytools, for each trait against hearing mode (amphibious hearing or in-air only), whether a taxon was aquatic (pinnipeds, otters, minks, polar bear) or terrestrial, and relative diving depth (above water, shallow depths, epipelagic, mesopelagic, bathypelagic). We used species mean trait values for these analyses.

**Estimating hearing of extinct pan-pinnipeds:** To estimate the hearing mode of extinct fossil pinnipeds, we used flexible discriminant analysis (FDA) of middle and inner ear morphometric data. The FDA models were assessed via confusion matrices and Jackknifing. The variation of morphometric data by hearing mode was visualised using a principal component analysis, and a multivariate analysis of variance (MANOVA) of the first 99% of PC scores was done against hearing mode. We then used an Equal Rates ancestral state estimation of hearing mode (in-air or amphibious hearing) for Caniformia using the R package phytools, which was the best supported model (assessed with AICc and log-likelihood values). We estimated the relative in-air low and high frequency hearing limits and ranges for all taxa using pre-existing equations ^50^, and relative underwater frequency hearing limits were estimated for amphibious hearing taxa from novel equations for this study (and also cetaceans for Figure 5). See Supplemental Information for details.

**Testing for evolutionary shifts:** We used BayesTraits ^97^ to investigate evolutionary rate shifts in morphological and auditory traits following the protocol in Brennan et al. ^98^, testing Brownian motion, variable rates, early burst, and Ornstein–Uhlenbeck models. We paid particular attention to evolutionary rate shifts that occurred near the origin of the evolution of amphibious hearing. See Supplemental Information for details.

**Morphological-auditory adaptive landscapes:** We ran a 3D geometric morphometric analysis of cochlear shape (which has functional correlates with hearing ranges) to construct adaptive landscapes, which we used to identify adaptive valleys and peaks in in-air and underwater hearing range. Cochlear meshes were landmarked using a modified version of the landmark protocol of Grohé *et al*. ^88^ and Taszus *et al.* ^14^, with 100 fixed equidistant landmarks (which preserves the homology of the cochlear curve) placed on the “dorsal” surface of the cochlear spiral in the program MorphoDig ^99^. A principal component analysis (PCA) of Procrustes 3D coordinates of cochlear shape was run using the R package geomorph to construct a morphospace. From this, we generated 80 theoretical cochlear morphologies (each represented as a set of 3D landmarks) in an 10 x 8 grid along PC1 and PC2. We used the estimated hearing range data (in octaves) for theoretical cochlear shapes to generate performance surfaces for in-air and underwater hearing ranges using the R package Morphoscape. These performance surfaces were used to calculate adaptive landscapes for hearing ability (in-air versus underwater) to assess adaptive tradeoffs for amphibious hearing. See Supplemental Information for details.

## Acknowledgements

We thank all museum collections staff who provided support and facilitated access, including Roberto Portela-Miguez, Phaedra Kokkini, Neil Adams (Natural History Museum, London), Jorge Velez-Juarbe (Natural History Museum of Los Angeles County), Nicholas Pyenson and Amanda Milhouse (National Museum of Natural History, Smithsonian Institution), Christian de Muizon and Géraldine Véron (Muséum national d’Histoire naturelle), Stephen Godfrey (Calvert Marine Museum), Romala Govender (Iziko Museums of South Africa), Anna Ďurišová (Slovak National Museum), Alan Tennyson (Museum of New Zealand Te Papa Tongarewa), Géraldine Garcia (PALEVOPRIM, University of Poitiers), Suzanne Jiquel, Bernard Marandat and Laurent Marivaux (Institut des Sciences de l’Evolution of Montpellier), and Nancy Simmons, Neil Duncan and Eileen Westwig (American Museum of Natural History, New York). We also are indebted to Brett Clark (NHM Imaging and Analysis center), Tautis Skorka (University of Southern California), Renaud Lebrun (Microtomography RX facility of Montpellier), Morgan Hill and Henry Towbin (Microscopy and Imaging Facility, AMNH of New York) for assistance with micro-CT scanning. Frances Paterson and the North Hobart Veterinary Hospital are thanked for assistance with medical-CT scanning of fresh seal heads. In addition, we thank the following researchers who kindly shared data: Natalia Rybczynski (scan of *Puijila*†), Anjali Goswami (scans of fossil phocids), and Stephen Godfrey (scan of *Leptophoca*†). Thanks to the NHM Macro Lab, Evans EvoMorph Lab, Marco Camaiti, Marc Jones, Hazel Richards, Erich Fitzgerald, Lindsey Koper, Joy Reidenberg, and Damian Haeusler for discussions that improved this manuscript. We thank Peter Trusler for providing silhouettes of pinnipeds.

## Data Availability Statement

All specimens used in this study are deposited and accessible in museum collections. 3D data scanned for this publication (and not previously published elsewhere) can be found in MorphoSource Project 000656781: SEAL: The Evolution of Auditory Adaptations for Aquatic Life in Pinnipeds. The data and code to replicate analyses, including landmarks and measurements, can be downloaded from Figshare, DOI: 10.6084/m9.figshare.31006549.

