## Supplemental Information, Supplemental Tables S1-S5, Supplemental Figures S1-S5, Supplemental Methods S1-S4 for "The origin and evolution of amphibious hearing in pinnipeds"

### The origin and evolution of amphibious hearing in pinnipeds: **Supplemental Information**

**Table S1. Specimen micro-CT scans used in this study from Methods.** Skeletally mature adult specimens were selected for scanning, indicated by skull suture fusion (including the fusion of the basisphenoid - basioccipital suture). When possible, the ear on the right side of the skull was scanned. Males and females were scanned (when possible) for pinnipeds, and for caniforms, musteloids, and ursids males were scanned when possible. Collection Abbreviations: American Museum of Natural History (AMNH), Calvert Marine Museum (CMM), Natural History Museum of Los Angeles County (LACM), Phyletic Museum Jena (MAM), Muséum national d'Histoire naturelle (MNHN), Monash University Zoology Collection (MZR), Natural History Museum London (NHMUK), Naturhistorisches Museum Basel (NMB), Museum of New Zealand Te Papa Tongarewa (NMNZ), Museums Victoria (NMV), Nunavut Fossil Vertebrate Collection, Canadian Museum of Nature (NUFV), Orange County's Paleontological Collection (OCPC), Iziko Museums of South Africa (SAM), San Diego Natural History Museum (SDSNH), Slovak National Museum (SNM), Sendai Science Museum (SSME), University of Montpellier collections (UM), University of Poitiers collections of extant and fossil vertebrates (UPPal), Smithsonian National Museum of Natural History (USNM), Natural History Museum Berlin (ZMB), Leipzig Zoo (Zoo).

| **Group** | **Taxon** | **Specimen number** | **Sex** | **Voxel size (um)** | **Source** |
|---|---|---|---|---|---|
| Canidae | *Vulpes vulpes* | MAM 2181 | ? | 20 | Taszus et al 2023 [1] |
| Canidae | *Vulpes vulpes* | MAM 2291 | Male | 20 | Taszus et al 2023 [1] |
| Canidae | *Vulpes vulpes* | MAM 751 | Female | 32.97 | Taszus et al 2023 [1] |
| Canidae | *Vulpes lagopus* | MAM 1007 | ? | 20 | Taszus et al 2023 [1] |
| Canidae | *Vulpes lagopus* | MAM 6838 | ? | 32.97 | Taszus et al 2023 [1] |
| Canidae | *Canis lupus* | MAM 3 | ? | 40 | Taszus et al 2023 [1] |
| Canidae | *Canis lupus* | MAM 5 | ? | 45 | Taszus et al 2023 [1] |
| Canidae | *Canis lupus* | MAM t | ? | 40.00 | Taszus et al 2023 [1] |
| Ursidae | *Ailuropoda melanoleuca* | NHMUK 39.3808 | Male | 29.10 | UKRI SEAL (this study) |
| Ursidae | *Ailuropoda melanoleuca* | MAM 37026 | ? | 62.28 | Taszus et al 2023 [1] |
| Ursidae | *Tremarctos ornatus* | NHMUK 27.11.1.70 | Male | 26.5 | UKRI SEAL (this study) |
| Ursidae | *Tremarctos ornatus* | Zoo specimen | ? | 63.77 | Taszus et al 2023 [1] |
| Ursidae | *Helarctos malayanus* | NHMUK 1938.11.30.69 | Male | 28.45 | UKRI SEAL (this study) |
| Ursidae | *Melursus ursinus* | NHMUK 32.5.7.9 | Male | 30.74 | UKRI SEAL (this study) |
| Ursidae | *Ursus arctos* | NHMUK 30.3.1.1 | Male | 26.86 | UKRI SEAL (this study) |
| Ursidae | *Ursus americanus* | NHMUK 61.1286 | Male | 23.38 | UKRI SEAL (this study) |
| Ursidae | *Ursus maritimus* | NHMUK 90.8.4.1 | Male | 31.08 | UKRI SEAL (this study) |
| Ursidae | *Ursus maritimus* | MAM 7323 | Female | 58.62 | Taszus et al 2023 [1] |
| Ursidae | *Ursus thibetanus* | NHMUK 26.10.8.41 | Male | 28.37 | UKRI SEAL (this study) |
| Musteloidea | *Lontra felina* | MNHN MO 1932-3019 | ? | 24.42 | Grohe et al 2016 [2] |
| Musteloidea | *Lontra canadensis* | AMNH M254476 | ? | 55.34 | Grohe et al 2016 [2] |
| Musteloidea | *Aonyx capensis* | NHMUK 1906.11.1.22 | Male | 17.02 | UKRI SEAL (this study) |
| Musteloidea | *Aonyx capensis* | MAM 6961 | ? | 30 | Taszus et al 2023 [1] |
| Musteloidea | *Aonyx cinerea* | MNHN MO 1982-162 | ? | 24.70 | Grohe et al 2016 [2] |
| Musteloidea | *Aonyx cinerea* | MAM 6958 | ? | 35 | Taszus et al 2023 [1] |
| Musteloidea | *Lutrogale perspicillata* | NHMUK 35.3.26.11 | Male | 12.89 | UKRI SEAL (this study) |
| Musteloidea | *Lutrogale perspicillata* | MAM 6950 | Male | 28 | Taszus et al 2023 [1] |
| Musteloidea | *Lutra lutra* | UPPal M02.5.005 | ? | 29.95 | Grohe et al 2016 [2] |
| Musteloidea | *Lutra lutra* | UM 009N | ? | 56.44 | Grohe et al 2016 [2] |
| Musteloidea | *Lutra lutra* | MAM 6956 | Female | 18.28 | Taszus et al 2023 [1] |
| Musteloidea | *Lutra lutra* | MAM 5136 | Male | 22.11 | Taszus et al 2023 [1] |
| Musteloidea | *Hydrictis maculicollis* | NHMUK 22.5.15.12 | Male | 10.50 | UKRI SEAL (this study) |
| Musteloidea | *Enhydra lutris* | MNHN MO 1935-124 | ? | 30.25 | Grohe et al 2016 [2] |
| Musteloidea | *Enhydra lutris* | AMNH 24186 | Male | 70.21 | Grohe et al 2016 [2] |
| Musteloidea | *Enhydra lutris* | ZMB 111 | ? | 34 | Taszus et al 2023 [1] |
| Musteloidea | *Pteronura brasiliensis* | MAM 6952 | ? | 33 | Taszus et al 2023 [1] |
| Musteloidea | *Mustela erminea* | MAM 6934 | Female | 14 | Taszus et al 2023 [1] |
| Musteloidea | *Mustela putorius* | UM 117N | ? | 24.83 | Grohe et al 2016 [2] |
| Musteloidea | *Mustela putorius* | MAM 894 | Male | 19 | Taszus et al 2023 [1] |
| Musteloidea | *Mustela putorius* | MAM 2326 | | 14.07 | Taszus et al 2023 [1] |
| Musteloidea | *Mustela lutreola* | UM 670N | ? | 27.36 | Grohe et al 2016 [2] |
| Musteloidea | *Mustela lutreola* | MAM 6909 | ? | 14 | Taszus et al 2023 [1] |
| Musteloidea | *Mustela itatsi* | NHMUK 5.5.30.8 | Male | 6.31 | UKRI SEAL (this study) |
| Musteloidea | *Mustela nivalis* | MNHN MO 1933-2153 | ? | 18.05 | Grohe et al 2016 [2] |

| | | | | | |
|---|---|---|---|---|---|
| Musteloidea | *Mustela nivalis* | MAM 6905 | ? | 15 | Taszus et al 2023 [1] |
| Musteloidea | *Mustela nivalis* | MAM 6921 | Male | 6.58 | Taszus et al 2023 [1] |
| Musteloidea | *Mustela kathiah* | NHMUK 33.4.1.249 | Male | 8.39 | UKRI SEAL (this study) |
| Musteloidea | *Mustela nudipes* | NHMUK 55.738 | Male | 9.11 | UKRI SEAL (this study) |
| Musteloidea | *Mustela nudipes* | MAM 6900 | ? | 14 | Taszus et al 2023 [1] |
| Musteloidea | *Mustela africana* | NHMUK 26.1.8.10 | Male | 7.53 | UKRI SEAL (this study) |
| Musteloidea | *Mustela frenata* | AMNH 60508 | ? | 31.41 | Grohe et al 2016 [2] |
| Musteloidea | *Neogale vision* | MNHN MO 1959-189 | ? | 31.21 | Grohe et al 2016 [2] |
| Musteloidea | *Neogale vision* | MAM 2155 | ? | 19 | Taszus et al 2023 [1] |
| Musteloidea | *Galictis cuja* | MNHN MO 1960-3811 | ? | 26.43 | Grohe et al 2016 [2] |
| Musteloidea | *Lyncodon patagonicus* | NHMUK 3.7.9.16 | Male | 9.14 | UKRI SEAL (this study) |
| Musteloidea | *Poecilogale albinucha* | NHMUK 68.252 | Male | 12.09 | UKRI SEAL (this study) |
| Musteloidea | *Ictonyx libycus* | NHMUK 12.6.12.55 | Male | 9.54 | UKRI SEAL (this study) |
| Musteloidea | *Ictonyx striatus* | MAM 6912 | Female | 14 | Taszus et al 2023 [1] |
| Musteloidea | *Vormela peregusna* | NHMUK 97.6.4.3 | Male | 8.10 | UKRI SEAL (this study) |
| Musteloidea | *Melogale moschata* | MNHN MO 1929-376 | ? | 24.72 | Grohe et al 2016 [2] |
| Musteloidea | *Melogale personata* | NHMUK 43.153 | Male | 16.23 | UKRI SEAL (this study) |
| Musteloidea | *Martes americana* | NHMUK 53.583 | Male | 15.48 | UKRI SEAL (this study) |
| Musteloidea | *Martes martes* | UPPal M02.5.019A | ? | 25.23 | Grohe et al 2016 [2] |
| Musteloidea | *Martes martes* | MAM 2084 | Male | 20 | Taszus et al 2023 [1] |
| Musteloidea | *Martes flavigula* | MAM 6867 | ? | 20 | Taszus et al 2023 [1] |
| Musteloidea | *Martes foina* | UPPal M02.5.017A | ? | 24.56 | Grohe et al 2016 [2] |
| Musteloidea | *Martes foina* | MAM 883 | Male | 35 | Taszus et al 2023 [1] |
| Musteloidea | *Martes foina* | MAM 2486 | ? | 17.54 | Taszus et al 2023 [1] |
| Musteloidea | *Gulo gulo* | MNHN MO 1873-39 | ? | 38.96 | Grohe et al 2016 [2] |
| Musteloidea | *Gulo gulo* | AMNH 182936 | ? | 73.81 | Grohe et al 2016 [2] |
| Musteloidea | *Gulo gulo* | MAM 6957 | ? | 34 | Taszus et al 2023 [1] |
| Musteloidea | *Pekania pennanti* | AMNH 121558 | ? | 50.15 | Grohe et al 2016 [2] |
| Musteloidea | *Eira barbara* | AMNH 32065 | ? | 52.72 | Grohe et al 2016 [2] |
| Musteloidea | *Eira barbara* | MAM 1001 | ? | 35 | Taszus et al 2023 [1] |
| Musteloidea | *Meles anakuma* | NHMUK 6.1.4.113 | Male | 11.50 | UKRI SEAL (this study) |
| Musteloidea | *Meles meles* | UPPal M0.2.5.021 | ? | 43.99 | Grohe et al 2016 [2] |
| Musteloidea | *Meles meles* | MAM 2918 | ? | 33 | Taszus et al 2023 [1] |
| Musteloidea | *Meles meles* | MAM 925 | Male | 27 | Taszus et al 2023 [1] |
| Musteloidea | *Meles meles* | MAM 2088 | Male | 25.58 | Taszus et al 2023 [1] |
| Musteloidea | *Arctonyx collaris* | MNHN MO 1962-153 | ? | 24.72 | Grohe et al 2016 [2] |
| Musteloidea | *Arctonyx collaris* | MAM 6615 | Female | 36 | Taszus et al 2023 [1] |
| Musteloidea | *Mellivora capensis* | MNHN MO 1893-6 | ? | 38.76 | Grohe et al 2016 [2] |
| Musteloidea | *Mellivora capensis* | MAM 6966 | ? | 24 | Taszus et al 2023 [1] |
| Musteloidea | *Taxidea taxus* | MNHN MO 1895-417 | ? | 27.21 | Grohe et al 2016 [2] |
| Musteloidea | *Taxidea taxus* | AMNH 120577 | ? | 82.33 | Grohe et al 2016 [2] |
| Musteloidea | *Taxidea taxus* | MAM 6962 | Male | 28 | Taszus et al 2023 [1] |
| Musteloidea | *Nasuella olivacea* | NHMUK 98.7.1.6 | Male | 9.85 | UKRI SEAL (this study) |
| Musteloidea | *Nasua nasua* | UM 141N | ? | 32.33 | Grohe et al 2016 [2] |
| Musteloidea | *Nasua nasua* | UPPal M02.5.024A | ? | 24.40 | Grohe et al 2016 [2] |
| Musteloidea | *Bassaricyon medius* | NHMUK 9.7.17.11 | Female | 9.97 | UKRI SEAL (this study) |
| Musteloidea | *Bassaricyon gabbii* | NHMUK 5.5.4.5 | Male | 9.69 | UKRI SEAL (this study) |
| Musteloidea | *Bassaricyon gabbi* | AMNH 47772 | ? | 62.5 | Grohe et al 2016 [2] |
| Musteloidea | *Procyon cancrivorous* | UM 002N | ? | 54.77 | Grohe et al 2016 [2] |
| Musteloidea | *Procyon lotor* | UM 091N | ? | 28.95 | Grohe et al 2016 [2] |
| Musteloidea | *Procyon lotor* | AMNH 24815 | ? | 88.14 | Grohe et al 2016 [2] |
| Musteloidea | *Procyon lotor* | MAM 2976 | ? | 20 | Taszus et al 2023 [1] |
| Musteloidea | *Procyon lotor* | MAM 8037 | Male | 20.11 | Taszus et al 2023 [1] |
| Musteloidea | *Bassariscus astutus* | AMNH 135964 | ? | 55.96 | Grohe et al 2016 [2] |
| Musteloidea | *Potos flavus* | UM 124N | ? | 73.14 | Grohe et al 2016 [2] |
| Musteloidea | *Potos flavus* | AMNH 239990 | ? | 72.21 | Grohe et al 2016 [2] |
| Musteloidea | *Ailurus fulgens* | MNHN MO 1963-358 | ? | 24.65 | Grohe et al 2016 [2] |
| Musteloidea | *Ailurus fulgens* | AMNH 185436 | ? | 94.42 | Grohe et al 2016 [2] |
| Musteloidea | *Spilogale putorius* | MNHN MO 1962-961 | ? | 72 | Grohe et al 2016 [2] |
| Musteloidea | *Spilogale putorius* | AMNH 35207 | ? | 57.68 | Grohe et al 2016 [2] |
| Musteloidea | *Mephitis mephitis* | MNHN MO 2005-655 | ? | 24.80 | Grohe et al 2016 [2] |

| | | | | | |
|---|---|---|---|---|---|
| Musteloidea | *Mephitis mephitis* | AMNH 172133 | ? | 58.33 | Grohe et al 2016 [2] |
| Musteloidea | *Mephitis mephitis* | MAM 901 | ? | 20 | Taszus et al 2023 [1] |
| Musteloidea | *Conepatus chinga* | NHMUK 9.12.1.16 | Male | 8.68 | UKRI SEAL (this study) |
| Musteloidea | *Mydaus javanensis* | AMNH 106635 | ? | 71.24 | Grohe et al 2016 [2] |
| Pinnipedia | *Arctocephalus pusillus* | NHMUK 1887.5.6.1 | Male | 31.37 | UKRI SEAL (this study) |
| Pinnipedia | *Arctocephalus pusillus* | NHMUK 1883.7.28.13 | Female | 24.70 | UKRI SEAL (this study) |
| Pinnipedia | *Arctocephalus gazella* | NHMUK 1960.8.4.4 | Male | 26.20 | UKRI SEAL (this study) |
| Pinnipedia | *Arctocephalus gazella* | NHMUK 1960.8.10.14 | Female | 24.36 | UKRI SEAL (this study) |
| Pinnipedia | *Arctocephalus townsendi* | AMNH 76844 | Male | 88.52 | Grohe et al 2018 [3] |
| Pinnipedia | *Arctocephalus philippii* | NHMUK 1883.11.8.1 | Male | 24.05 | UKRI SEAL (this study) |
| Pinnipedia | *Arctocephalus galapagoensis* | NHMUK ZD 1991.1 | Male | 24.05 | UKRI SEAL (this study) |
| Pinnipedia | *Arctocephalus galapagoensis* | NHMUK ZD 1991.2 | Female | 21.23 | UKRI SEAL (this study) |
| Pinnipedia | *Arctocephalus forsteri* | NHMUK 1968.9.26.6 | Male | 27.07 | UKRI SEAL (this study) |
| Pinnipedia | *Arctocephalus forsteri* | NHMUK 1968.9.26.5 | Female | 23.59 | UKRI SEAL (this study) |
| Pinnipedia | *Arctocephalus tropicalis* | NHMUK 1957.4.23.11 | Male | 20.83 | UKRI SEAL (this study) |
| Pinnipedia | *Arctocephalus tropicalis* | NHMUK 1955.3.14.5 | Female | 23.06 | UKRI SEAL (this study) |
| Pinnipedia | *Arctocephalus australis* | NHMUK 1984.973 | Male | 24.05 | UKRI SEAL (this study) |
| Pinnipedia | *Arctocephalus australis* | NHMUK 1984.919 | Female | 24.36 | UKRI SEAL (this study) |
| Pinnipedia | *Callorhinus ursinus* | NHMUK 1893.1.28.2 | Male | 27.07 | UKRI SEAL (this study) |
| Pinnipedia | *Callorhinus ursinus* | NHMUK 1960.5.2.2 | Female | 26.36 | UKRI SEAL (this study) |
| Pinnipedia | *Callorhinus ursinus* | ZMB 5628 | ? | 31 | Taszus et al 2023 [1] |
| Pinnipedia | *Callorhinus ursinus* | ZMB 5627 | ? | 46 | Taszus et al 2023 [1] |
| Pinnipedia | *Neophoca cinerea* | NHMUK 1897.10.10.5 | Male | 31.50 | UKRI SEAL (this study) |
| Pinnipedia | *Neophoca cinerea* | NHMUK 1968.9.26.27 | Female | 32.22 | UKRI SEAL (this study) |
| Pinnipedia | *Eumetopias jubatus* | NHMUK 1950.3.29.11 | Male | 35.78 | UKRI SEAL (this study) |
| Pinnipedia | *Eumetopias jubatus* | MAM 72815 | ? | 76.94 | Taszus et al 2023 [1] |
| Pinnipedia | *Otaria flavescens/byronia* | NHMUK 1939.1.21.166 | Male | 31.01 | UKRI SEAL (this study) |
| Pinnipedia | *Otaria flavescens/byronia* | NHMUK 84.991 | Female | 28.62 | UKRI SEAL (this study) |
| Pinnipedia | *Phocarctos hookeri* | NHMUK 1876.2.16.9 | Male | 26.66 | UKRI SEAL (this study) |
| Pinnipedia | *Phocarctos hookeri* | NHMUK 1908.2.20.51 | Female | 26.36 | UKRI SEAL (this study) |
| Pinnipedia | *Zalophus californianus* | NHMUK 1968.6.10.1 | Male | 27.07 | UKRI SEAL (this study) |
| Pinnipedia | *Zalophus californianus* | NHMUK 1903.10.11.4 | Female | 26.36 | UKRI SEAL (this study) |
| Pinnipedia | *Zalophus californianus* | MZRC 105 | ? | 41.26 | UKRI SEAL (this study) |
| Pinnipedia | *Zalophus californianus* | MAM 8274 | ? | 30 | Taszus et al 2023 [1] |
| Pinnipedia | *Zalophus japonicus†* | NHMUK 1873.3.12.1 | Male | 30.50 | UKRI SEAL (this study) |
| Pinnipedia | *Zalophus wollebaeki* | AMNH 63946 | Male | 86.78 | Grohe et al 2018 [3] |
| Pinnipedia | *Odobenus rosmarus* | NHMUK 39.4474B | ? | 52.90 | UKRI SEAL (this study) |
| Pinnipedia | *Odobenus rosmarus* | NHMUK 1855.11.26.37 | Female | 90.34 | UKRI SEAL (this study) |
| Pinnipedia | *Odobenus rosmarus* | MAM 648 | ? | 100.75 | Taszus et al 2023 [1] |
| Pinnipedia | *Erignathus barbatus* | AMNH 98 | ? | 92.63 | Grohe et al 2018 [3] |
| Pinnipedia | *Erignathus barbatus* | NHMUK 1937.10.23.9 | Female | 23.45 | UKRI SEAL (this study) |
| Pinnipedia | *Cystophora cristata* | NHMUK 1890.8.1.4 | Female | 36.78 | UKRI SEAL (this study) |
| Pinnipedia | *Cystophora cristata* | NHMUK 332h | Male | 41.95 | UKRI SEAL (this study) |
| Pinnipedia | *Phoca vitulina* | AMNH 100 | ? | 85.53 | Grohe et al 2018 [3] |
| Pinnipedia | *Phoca vitulina* | NHMUK 1868.3.21.1 | Male | 21.29 | UKRI SEAL (this study) |
| Pinnipedia | *Phoca vitulina* | NHMUK 1928.9.1.2 | Female | 24.66 | UKRI SEAL (this study) |
| Pinnipedia | *Phoca vitulina* | MAM 2982 | ? | 45.80 | Taszus et al 2023 [1] |
| Pinnipedia | *Phoca largha* | NHMUK 1965.7.19.11 | Female | 26.36 | UKRI SEAL (this study) |
| Pinnipedia | *Phoca largha* | NHMUK 1965.7.19.13 | Male | 23.45 | UKRI SEAL (this study) |
| Pinnipedia | *Pusa hispida* | NHMUK 1937.10.23.1 | Male | 21.82 | UKRI SEAL (this study) |
| Pinnipedia | *Pusa hispida* | NHMUK 1938.12.3.1 | Female | 22.75 | UKRI SEAL (this study) |
| Pinnipedia | *Pusa hispida* | MAM 1270 | ? | 45.80 | Taszus et al 2023 [1] |
| Pinnipedia | *Pusa sibirica* | NHMUK 1963.7.19.9 | Male | 21.10 | UKRI SEAL (this study) |
| Pinnipedia | *Pusa sibirica* | NHMUK 1965.9.6.1 | Female | 19.23 | UKRI SEAL (this study) |
| Pinnipedia | *Pusa caspica* | NHMUK 1965.7.19.1 | Male | 22.34 | UKRI SEAL (this study) |
| Pinnipedia | *Pusa caspica* | NHMUK 1965.7.19.2 | Female | 18.04 | UKRI SEAL (this study) |
| Pinnipedia | *Halichoerus grypus* | NHMUK 1961.1.23.3 | Female | 36.78 | UKRI SEAL (this study) |
| Pinnipedia | *Halichoerus grypus* | NHMUK 1961.5.18.12 | Male | 36.78 | UKRI SEAL (this study) |
| Pinnipedia | *Halichoerus grypus* | MAM 2387 | ? | 37 | Taszus et al 2023 [1] |
| Pinnipedia | *Pagophilus groenlandicus* | NHMUK 1938.12.10.1 | Female | 29.65 | UKRI SEAL (this study) |
| Pinnipedia | *Pagophilus groenlandicus* | NHMUK 1963.7.19.4 | Male | 20.41 | UKRI SEAL (this study) |

| | | | | | |
|---|---|---|---|---|---|
| Pinnipedia | *Pagophilus groenlandicus* | ZMB 56776 | ? | 44 | Taszus et al 2023 [1] |
| Pinnipedia | *Pagophilus groenlandicus* | ZMB 32569 | ? | 37 | Taszus et al 2023 [1] |
| Pinnipedia | *Pagophilus groenlandicus* | MAM 627 | ? | 45.80 | Taszus et al 2023 [1] |
| Pinnipedia | *Histriophoca fasciata* | NHMUK 1965.7.19.8 | Male | 29.67 | UKRI SEAL (this study) |
| Pinnipedia | *Histriophoca fasciata* | NHMUK 1965.7.19.9 | Female | 29.67 | UKRI SEAL (this study) |
| Pinnipedia | *Monachus monachus* | NHMUK 1863.4.1.1 | Male | 117.65 | MSCA ECHO (this study) |
| Pinnipedia | *Monachus monachus* | NHMUK 1892.10.4.1 | ? | 127.59 | MSCA ECHO (this study) |
| Pinnipedia | *Monachus monachus* | NHMUK 1894.7.27.1 | Male | 127.59 | MSCA ECHO (this study) |
| Pinnipedia | *Monachus monachus* | NHMUK 1894.7.27.2 | Female | 127.59 | MSCA ECHO (this study) |
| Pinnipedia | *Monachus monachus* | NHMUK 1934.8.5.4 | ? | 127.59 | MSCA ECHO (this study) |
| Pinnipedia | *Neomonachus tropicalis†* | NHMUK 1889.11.5.1 | Male | 127.59 | MSCA ECHO (this study) |
| Pinnipedia | *Neomonachus schauinslandi* | NHMUK 1958.11.26.1 | Female | 118.03 | MSCA ECHO (this study) |
| Pinnipedia | *Mirounga leonina* | NHMUK 1933.8.12.1 | Male | 49.23 | UKRI SEAL (this study) |
| Pinnipedia | *Mirounga leonina* | NHMUK 1954.5.20.8 | Female | 44.19 | UKRI SEAL (this study) |
| Pinnipedia | *Mirounga leonina* | AMNH 48161 | Male | 141.73 | Grohe et al 2018 [3] |
| Pinnipedia | *Mirounga angustirostris* | AMNH 32677 | Female | 113.77 | Grohe et al 2018 [3] |
| Pinnipedia | *Mirounga angustirostris* | MAM 4196 | Female | 73.27 | Taszus et al 2023 [1] |
| Pinnipedia | *Lobodon carcinophaga* | NHMUK 1940.4.6.93 | Male | 28.55 | UKRI SEAL (this study) |
| Pinnipedia | *Lobodon carcinophaga* | NHMUK 1959.12.8.9 | Female | 33.01 | UKRI SEAL (this study) |
| Pinnipedia | *Lobodon carcinophaga* | AMNH 88498 | Male | 90.65 | Grohe et al 2018 [3] |
| Pinnipedia | *Hydrurga leptonyx* | NHMUK 1939.2.11.20 | Male | 31.78 | UKRI SEAL (this study) |
| Pinnipedia | *Hydrurga leptonyx* | NHMUK 1940.4.6.142 | Female | 39.88 | UKRI SEAL (this study) |
| Pinnipedia | *Hydrurga leptonyx* | AMNH 34920 | ? | 124.71 | Grohe et al 2018 [3] |
| Pinnipedia | *Leptonychotes weddellii* | NHMUK 1940.4.6.83 | Male | 29.35 | UKRI SEAL (this study) |
| Pinnipedia | *Leptonychotes weddellii* | NHMUK 1940.4.6.20 | Female | 35.52 | UKRI SEAL (this study) |
| Pinnipedia | *Leptonychotes weddellii* | AMNH 88548 | ? | 99.91 | Grohe et al 2018 [3] |
| Pinnipedia | *Leptonychotes weddellii* | MAM 2133 | ? | 54 | Taszus et al 2023 [1] |
| Pinnipedia | *Ommatophoca rossi* | NHMUK 1949.2.3.1 | ? | 101.53 | MSCA ECHO (this study) |
| Pinnipedia | *Ommatophoca rossi* | NHMUK 1901.1.4.12 | ? | 114.65 | MSCA ECHO (this study) |
| Pinnipedia | *Homiphoca capensis†* | SAM-PQ-48759 | ? | 62.63 | UKRI SEAL (this study) |
| Pinnipedia | *Hadrokirus martini†* | MNHN-F-SAS 1627a | ? | 49.52 | Daniela Sanfelice (this study) |
| Pinnipedia | *Acrophoca longirostris†* | MNHN-F-SAS 1273 | ? | 48.62 | Daniela Sanfelice (this study) |
| Pinnipedia | *Piscophoca pacifica†* | MNHN-F-SAS 564 | ? | 52.58 | Daniela Sanfelice (this study) |
| Pinnipedia | *Eomonachus belegaerensis†* | NMNZ.S.056361 | ? | 15.00 | Felix Marx (this study) |
| Pinnipedia | *"Leptophoca lenis"†* | CMM-V-2021 | ? | 66.63 | Stephen Godfrey |
| Pinnipedia | *Devinophoca claytoni†* | SNM Z14523 | ? | 60.00 | Martin Sabol (this study) |
| Pinnipedia | *Sarcodectes magnus†* | USNM PAL 181601 | ? | 54.75 | Stephanie Palmer (this study) |
| Pinnipedia | *Potamotherium vallentoni†* | NHMUK PV M11718 | ? | 18.75 | UKRI SEAL (this study) |
| Pinnipedia | *Puijila darwini†* | NUFV-405 | ? | 30 | Natalia Rybczynski + Ryan Paterson |
| Pinnipedia | *Enaliarctos mealsi†* | LACM 4321 | ? | 60.57 // 32.38 | UKRI SEAL (this study) |
| Pinnipedia | *Enaliarctos mitchelli†* | USNM PAL 175637 | ? | 92.90 | Stephanie Palmer (this study) |
| Pinnipedia | *Pteronarctos goedertae†* | LACM 123883 | ? | 55.58 | UKRI SEAL (this study) |
| Pinnipedia | *Pinnarctidion* sp.*†* | LACM 127974 | ? | 49.68 | UKRI SEAL (this study) |
| Pinnipedia | *Pinnarctidion* sp. c.f. *iverseni†* | USNM PAL 314312 | ? | 44.64 | Stephanie Palmer (this study) |
| Pinnipedia | *Allodesmus kernensis†* | LACM 4320 | ? | 102.53 | UKRI SEAL (this study) |
| Pinnipedia | *Desmatophoca brachycephala†* | LACM 120199 | ? | 50.8 | UKRI SEAL (this study) |
| Pinnipedia | *Prototaria planicephala†* | SSME13317 | ? | 37.43 | Naoki Kohno (this study) |
| Pinnipedia | *Neotherium mirum†* | LACM 52172 | ? | 46.15 | UKRI SEAL (this study) |
| Pinnipedia | *Pontolis barroni†* | LACM 123283 | ? | 84.56 | UKRI SEAL (this study) |
| Pinnipedia | *Pithanotaria starri†* | LACM 122619 | ? | 34.00 | UKRI SEAL (this study) |
| Pinnipedia | *Callorhinus* sp.*†* | OCPC 1894 | ? | 43.47 | UKRI SEAL (this study) |
| Cetacea | *Balaenoptera physalus* | NHMUK 1998.30.2 | ? | 40.05 | Park et al 2023 [4] |
| Cetacea | *Caperea marginata* | NMV C28531 | ? | 23.6 | Park et al 2017 [5] |
| Cetacea | *Delphinapterus leucas* | NMBCIII 1086 | ? | 40 | Costeur et al 2018 [6] |
| Cetacea | *Eschrichtius robustus* | SDNHM 24316 | ? | 29.3 | Ekdale & Racicot 2015 [7] |
| Cetacea | *Eubalaena australis* | NHMUK 1873.3.3.1 | ? | 21.05 | Park et al 2023 [4] |
| Cetacea | *Inia geoffrensis* | NMB 7167 | ? | 30 | Aguirre-Fernandez et al 2017 [8] |
| Cetacea | *Kogia breviceps* | NHMC 24976 | ? | 33.07 | Park et al 2016 [9] |
| Cetacea | *Lipotes vexillifer* | AMNH 57333 | ? | 30.7 | Costeur et al 2018 [6] |
| Cetacea | *Phocoena phocoena* | NMNH 571892 | ? | 42 | Racicot et al 2016 [10] |
| Cetacea | *Physeter macrocephalus* | NHMUK CE893.2 | ? | 42.14 | Park et al 2019 [11] |

| | | | | | |
|---|---|---|---|---|---|
| Cetacea | *Platanista gangetica* | NMV C27417.2 | ? | 41.5 | Park et al 2016 [9] |
| Cetacea | *Pontoporia blainvillei* | MNHN 1928.168 | ? | 38 | Costeur et al 2018 [6] |
| Cetacea | *Tasmacetus shepherdi* | NMV C37967.6.2 | ? | 58.28 | Park et al 2016 [9] |
| Cetacea | *Tursiops truncatus* | SDSNH 21212 | ? | 39.1 | Ekdale 2013 [12] |

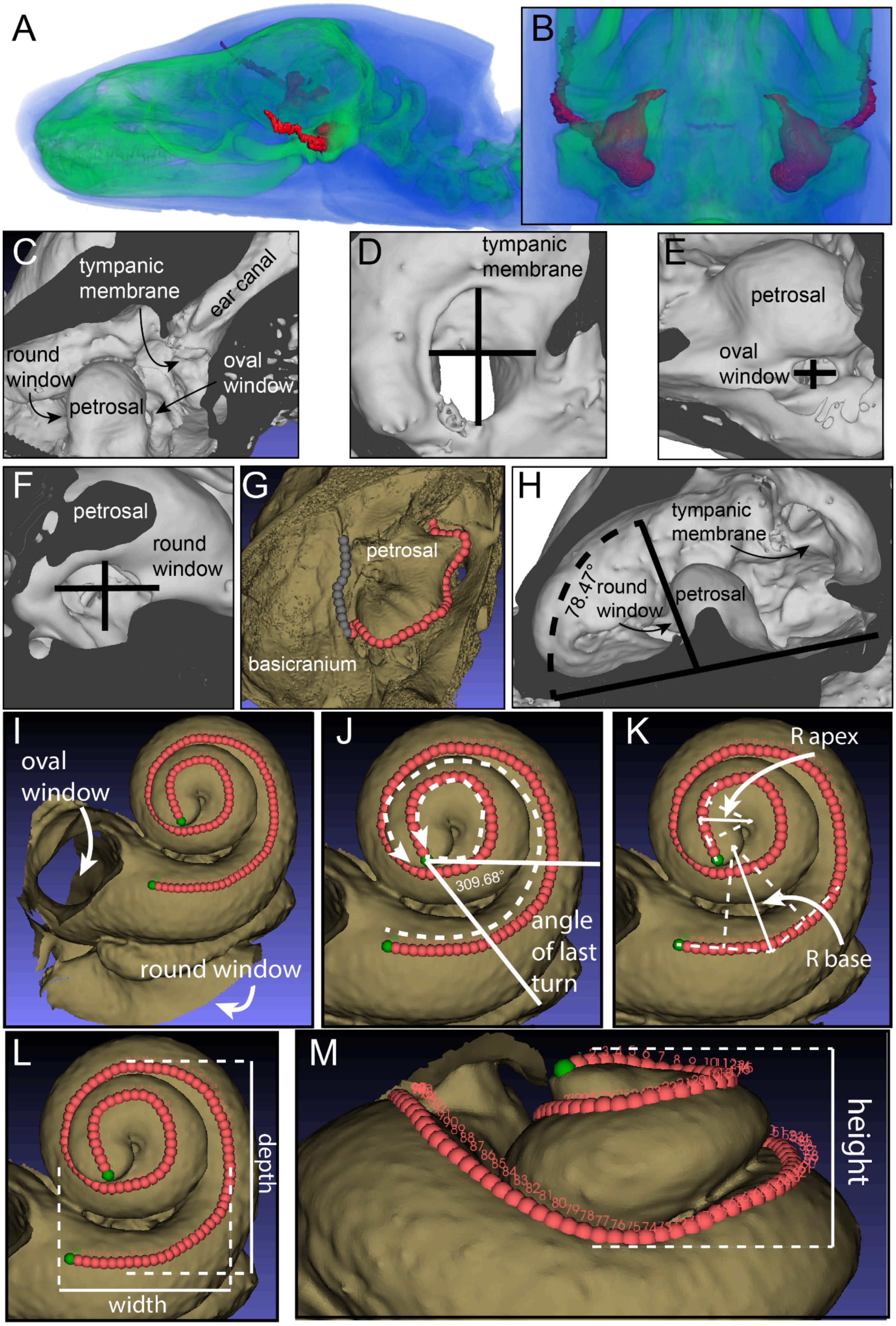


**Figure S1. Specimen measurements as described in Methods**. (A) oblique lateral view and (B) ventral view of TMAG A11092 with cavernous sinus segmented in red and bone in green. (C) oblique cross section showing position of auditory morphology. (D) tympanic membrane measurements. (E) oval window measurements. (F) round window measurements. (G) landmarks of the petrosal border (dorsal view, landmarks contacting basicranium in grey). (H) angle of the round window to the petrosal axis (oblique medial view). (I) 100 equidistant landmarks on the "dorsal" surface of the cochlea. (J) turn number, with incomplete turn calculated from angle measurement. (K) radii ratio of outer turn and inner turn of cochlea. (L) width and depth, and (M) height of cochlea used to calculate height-width ratio and geometric mean. Specimen shown is *Hydrurga leptonyx* NHMUK 1940.4.6.142, not to scale.

**Table S2. Categorical assignments from PGLS analysis in Methods.** Species assignments for hearing mode, dive depth, and habitat, and sources of predictor variable data. Categories are defined as follows: In-Air Hearing occurs entirely above water and in-air, where airborne soundwaves are received and processed. Underwater Hearing, where waterborne soundwaves are received and processed entirely underwater. Amphibious Hearing, where hearing can occur either in-air or underwater, and hearing ability underwater is as clear and effective as in-air hearing. Terrestrial, where animals live entirely on land, and therefore live surrounded by air. Aquatic, where animals live in some form around, in, or underneath water. Above water, animals that rarely dive underwater. Shallow diving, an inconsistent diving behaviour below 10 meters depth, and/or very infrequent. Defines freshwater divers such as otters and minks, as well as shallow ocean divers such as the sea otter and polar bear. Epipelagic, the uppermost layer of the ocean, or sunlight zone, between 0 and 200 meters depth. This layer of the ocean represents the extent light penetrates. Mesopelagic, the twilight zone, between 200-1000 meters depth. Very little light penetrates this layer of the ocean. Bathypelagic, the midnight zone, between 1000-4000 meters depth. No light reaches this depth, it is a consistent temperature and salinity, and has hydrostatic temperature over 100 atmospheres.

| **Taxon** | **Hearing mode** | **Diving Depth** | **Habitat** | **Reference** |
|---|---|---|---|---|
| *Ailuropoda melanoleuca* | In-air only | Above water | Terrestrial | Nowak 2003 [13] |
| *Ailurus fulgens* | In-air only | Above water | Terrestrial | Nowak 2003 [13] |
| *Aonyx capensis* | In-air only | Shallow | Aquatic | Thewissen and Nummela 2008 [14] |
| *Aonyx cinerea* | In-air only | Shallow | Aquatic | Thewissen and Nummela 2008 [14] |
| *Arctocephalus australis* | Amphibious | Epipelagic | Aquatic | Schreer and Kovacs 1997; Houser 2025; Reichmuth et al 2013;Thewissen and Nummela 2008 [15–17] |
| *Arctocephalus forsteri* | Amphibious | Mesopelagic | Aquatic | Page et al. 2005; Houser 2025; Reichmuth et al 2013; Thewissen and Nummela 2008 [15,16,18] |
| *Arctocephalus galapagoensis* | Amphibious | Epipelagic | Aquatic | Horning and Trillmich 1997; Houser 2025; Reichmuth et al 2013; Thewissen and Nummela 2008 [14–16,19] |
| *Arctocephalus gazella* | Amphibious | Mesopelagic | Aquatic | Schreer and Kovacs 1997; Lea et al 2002; Houser 2025; Reichmuth et al 2013; Thewissen and Nummela 2008 [14–17,20] |
| *Arctocephalus philippii* | Amphibious | Epipelagic | Aquatic | Francis et al. 1998; Houser 2025; Reichmuth et al 2013; Thewissen and Nummela 2008 [14–16,21] |
| *Arctocephalus pusillus* | Amphibious | Mesopelagic | Aquatic | Schreer and Kovacs 1997; Houser 2025; Reichmuth et al 2013; Thewissen and Nummela 2008 [14–17] |
| *Arctocephalus townsendi* | Amphibious | Epipelagic | Aquatic | Lander et al. 2000; Houser 2025; Reichmuth et al 2013; Thewissen and Nummela 2008 [14–16,22] |
| *Arctocephalus tropicalis* | Amphibious | Mesopelagic | Aquatic | Georges et al. 2000; Houser 2025; Reichmuth et al 2013; Thewissen and Nummela 2008 [14–16,23] |
| *Arctonyx collaris* | In-air only | Abovewater | Terrestrial | Nowak 2003 [13] |
| *Balaenoptera physalus* | Underwater only | NA | NA | Nowak 2003; Thewissen and Nummela 2008 [13,14] |
| *Bassaricyon gabbii* | In-air only | Above water | Terrestrial | Nowak 2003 [13] |
| *Bassaricyon medius* | In-air only | Above water | Terrestrial | Nowak 2003 [13] |
| *Bassariscus astutus* | In-air only | Above water | Terrestrial | Nowak 2003 [13] |
| *Callorhinus ursinus* | Amphibious | Mesopelagic | Aquatic | Gentry 2009; Ponganis et al. 1992; Houser 2025; Reichmuth et al 2013; Thewissen and Nummela 2008 [14–16,24,25] |

| | | | | |
|---|---|---|---|---|
| *Canis lupus* | In-air only | Above water | Terrestrial | Nowak 2003 [13] |
| *Caperea marginata* | Underwater only | NA | NA | Nowak 2003; Thewissen and Nummela 2008 [13,14] |
| *Conepatus chinga* | In-air only | Above water | Terrestrial | Nowak 2003 [13] |
| *Cystophora cristata* | Amphibious | Bathypelagic | Aquatic | Folkow and Blix 1999; Houser 2025; Reichmuth et al 2013; Thewissen and Nummela 2008 [14–16,26] |
| *Delphinapterus leucas* | Underwater only | NA | NA | Nowak 2003; Thewissen and Nummela 2008 [13,14] |
| *Eira barbara* | In-air only | Above water | Terrestrial | Nowak 2003 [13] |
| *Enhydra lutris* | In-air only | Shallow | Aquatic | Thewissen and Nummela 2008 [14] |
| *Erignathus barbatus* | Amphibious | Mesopelagic | Aquatic | Schreer and Kovacs 1997, IUCN; Houser 2025; Reichmuth et al 2013; Thewissen and Nummela 2008 [14–17] |
| *Eschrichtius robustus* | Underwater only | NA | NA | Nowak 2003; Thewissen and Nummela 2008 [13,14] |
| *Eubalaena australis* | Underwater only | NA | NA | Nowak 2003; Thewissen and Nummela 2008 [13,14] |
| *Eumetopias jubatus* | Amphibious | Mesopelagic | Aquatic | Schreer and Kovacs 1997; Pitcher et al 2005; Houser 2025; Reichmuth et al 2013; Thewissen and Nummela 2008 [14–17] |
| *Galictis cuja* | In-air only | Above water | Terrestrial | Nowak 2003 [13] |
| *Gulo gulo* | In-air only | Above water | Terrestrial | Nowak 2003 [13] |
| *Halichoerus grypus* | Amphibious | Mesopelagic | Aquatic | Beck et al. 2003; IUCN; Houser 2025; Reichmuth et al 2013; Thewissen and Nummela 2008 [14–16,27] |
| *Helarctos malayanus* | In-air only | Above water | Terrestrial | Nowak 2003 [13] |
| *Histriophoca fasciata* | Amphibious | Mesopelagic | Aquatic | IUCN; Houser 2025; Reichmuth et al 2013; Thewissen and Nummela 2008 [14–16] |
| *Hydrictis maculicollis* | In-air only | Shallow | Aquatic | Thewissen and Nummela 2008 [14] |
| *Hydrurga leptonyx* | Amphibious | Bathypelagic | Aquatic | Kienle et al 2022; Houser 2025; Reichmuth et al 2013; Thewissen and Nummela 2008 [14–16,28] |
| *Ictonyx libycus* | In-air only | Above water | Terrestrial | Nowak 2003 [13] |
| *Ictonyx striatus* | In-air only | Above water | Terrestrial | Nowak 2003 [13] |
| *Inia geoffrensis* | Underwater only | NA | NA | Nowak 2003; Thewissen and Nummela 2008 [13,14] |
| *Kogia breviceps* | Underwater only | NA | NA | Nowak 2003; Thewissen and Nummela 2008 [13,14] |
| *Leptonychotes weddellii* | Amphibious | Mesopelagic | Aquatic | Heerah et al. 2013; Houser 2025; Reichmuth et al 2013; Thewissen and Nummela 2008 [14–16,29] |

| | | | | |
|---|---|---|---|---|
| *Lipotes vexillifer* | Underwater only | NA | NA | Nowak 2003; Thewissen and Nummela 2008 [13,14] |
| *Lobodon carcinophaga* | Amphibious | Mesopelagic | Aquatic | Nachtsheim et al. 2017; Harcourt 2001; Houser 2025; Reichmuth et al 2013; Thewissen and Nummela 2008 [14–16,30] |
| *Lontra canadensis* | In-air only | Shallow | Aquatic | Thewissen and Nummela 2008 [14] |
| *Lontra felina* | In-air only | Shallow | Aquatic | Thewissen and Nummela 2008 [14] |
| *Lutra lutra* | In-air only | Shallow | Aquatic | Thewissen and Nummela 2008 [14] |
| *Lutrogale perspicillata* | In-air only | Shallow | Aquatic | Thewissen and Nummela 2008 [14] |
| *Lyncodon patagonicus* | In-air only | Above water | Terrestrial | Nowak 2003 [13] |
| *Martes americana* | In-air only | Above water | Terrestrial | Nowak 2003 [13] |
| *Martes flavigula* | In-air only | Above water | Terrestrial | Nowak 2003 [13] |
| *Martes foina* | In-air only | Above water | Terrestrial | Nowak 2003 [13] |
| *Martes martes* | In-air only | Above water | Terrestrial | Nowak 2003 [13] |
| *Meles anakuma* | In-air only | Above water | Terrestrial | Nowak 2003 [13] |
| *Meles meles* | In-air only | Above water | Terrestrial | Nowak 2003 [13] |
| *Mellivora capensis* | In-air only | Above water | Terrestrial | Nowak 2003 [13] |
| *Melogale moschata* | In-air only | Above water | Terrestrial | Nowak 2003 [13] |
| *Melogale personata* | In-air only | Above water | Terrestrial | Nowak 2003 [13] |
| *Melursus ursinus* | In-air only | Above water | Terrestrial | Nowak 2003 [13] |
| *Mephitis mephitis* | In-air only | Above water | Terrestrial | Nowak 2003 [13] |
| *Mirounga angustirostris* | Amphibious | Bathypelagic | Aquatic | Schreer and Kovacs 1997, IUCN; Houser 2025; Reichmuth et al 2013; Thewissen and Nummela 2008 [14–17] |
| *Mirounga leonina* | Amphibious | Bathypelagic | Aquatic | McIntyre et al. 2010; Houser 2025; Reichmuth et al 2013; Thewissen and Nummela 2008 [14–16,31] |
| *Monachus monachus* | Amphibious | Mesopelagic | Aquatic | IUCN; Houser 2025; Reichmuth et al 2013; Thewissen and Nummela 2008 [14–16] |
| *Mustela africana* | In-air only | Above water | Terrestrial | Nowak 2003 [13] |
| *Mustela erminea* | In-air only | Above water | Terrestrial | Nowak 2003 [13] |
| *Mustela frenata* | In-air only | Above water | Terrestrial | Nowak 2003 [13] |
| *Mustela itatsi* | In-air only | Above water | Terrestrial | Nowak 2003 [13] |
| *Mustela kathiah* | In-air only | Above water | Terrestrial | Nowak 2003 [13] |
| *Mustela lutreola* | In-air only | Shallow | Aquatic | Nowak 2003; Thewissen and Nummela 2008 [13,14] |

| | | | | |
|---|---|---|---|---|
| *Mustela nivalis* | In-air only | Above water | Terrestrial | Nowak 2003 [13] |
| *Mustela nudipes* | In-air only | Above water | Terrestrial | Nowak 2003 [13] |
| *Mustela putorius* | In-air only | Above water | Terrestrial | Nowak 2003 [13] |
| *Mydaus javanensis* | In-air only | Above water | Terrestrial | Nowak 2003 [13] |
| *Nasua nasua* | In-air only | Above water | Terrestrial | Nowak 2003 [13] |
| *Nasuella olivacea* | In-air only | Above water | Terrestrial | Nowak 2003 [13] |
| *Neogale vision* | In-air only | Shallow | Aquatic | Nowak 2003; Thewissen and Nummela 2008 [13,14] |
| *Neomonachus schauinslandi* | Amphibious | Mesopelagic | Aquatic | Parrish et al. 2002; Nowak 1999; IUCN; Houser 2025; Reichmuth et al 2013; Thewissen and Nummela 2008 [13–16,32] |
| *Neophoca cinerea* | Amphibious | Epipelagic | Aquatic | Costa and Gales 2003; Houser 2025; Reichmuth et al 2013; Thewissen and Nummela 2008 [14–16,33] |
| *Odobenus rosmarus* | Amphibious | Mesopelagic | Aquatic | Garde et al 2018; Houser 2025; Reichmuth et al 2013; Thewissen and Nummela 2008 [14–16,34] |
| *Ommatophoca rossii* | Amphibious | Mesopelagic | Aquatic | Blix and Nordoy 2007; Houser 2025; Reichmuth et al 2013; Thewissen and Nummela 2008 [14–16,35] |
| *Otaria byronia* | Amphibious | Epipelagic | Aquatic | Hückstädt et al. 2016; Houser 2025; Reichmuth et al 2013; Thewissen and Nummela 2008 [14–16,36] |
| *Pagophilus groenlandicus* | Amphibious | Mesopelagic | Aquatic | Folkow et al. 2004; Houser 2025; Reichmuth et al 2013; Thewissen and Nummela 2008 [14–16,37] |
| *Pekania pennanti* | In-air only | Above water | Terrestrial | Nowak 2003 [13] |
| *Phoca largha* | Amphibious | Epipelagic | Aquatic | Schreer and Kovacs 1997; Houser 2025; Reichmuth et al 2013; Thewissen and Nummela 2008 [14–17] |
| *Phoca vitulina* | Amphibious | Mesopelagic | Aquatic | Rosing-Asvid et al. 2020; Houser 2025; Reichmuth et al 2013; Thewissen and Nummela 2008 [14–16,38] |
| *Phocarctos hookeri* | Amphibious | Mesopelagic | Aquatic | Chilvers et al. 2005; Houser 2025; Reichmuth et al 2013; Thewissen and Nummela 2008 [14–16,39] |
| *Phocoena phocoena* | Underwater only | NA | NA | Nowak 2003; Thewissen and Nummela 2008 [13,14] |
| *Physeter macrocephalus* | Underwater only | NA | NA | Nowak 2003; Thewissen and Nummela 2008 [13,14] |
| *Platanista gangetica* | Underwater only | NA | NA | Nowak 2003; Thewissen and Nummela 2008 [13,14] |
| *Poecilogale albinucha* | In-air only | Above water | Terrestrial | Nowak 2003 [13] |
| *Pontoporia blainvillei* | Underwater only | NA | NA | Nowak 2003; Thewissen and Nummela 2008 [13,14] |
| *Potos flavus* | In-air only | Above water | Terrestrial | Nowak 2003 [13] |
| *Procyon cancrivorus* | In-air only | Above water | Terrestrial | Nowak 2003 [13] |
| *Procyon lotor* | In-air only | Above water | Terrestrial | Nowak 2003 [13] |

| | | | | |
|---|---|---|---|---|
| *Pteronura brasiliensis* | In-air only | Shallow | Aquatic | Thewissen and Nummela 2008 [14] |
| *Pusa caspica* | Amphibious | Epipelagic | Aquatic | Dmitrieva et al 2016; Houser 2025; Reichmuth et al 2013; Thewissen and Nummela 2008 [14–16,40] |
| *Pusa hispida* | Amphibious | Mesopelagic | Aquatic | Schreer and Kovacs 1997; IUCN; Houser 2025; Reichmuth et al 2013; Thewissen and Nummela 2008 [14–17] |
| *Pusa sibirica* | Amphibious | Mesopelagic | Aquatic | Stewart et al. 1996; Schreer and Kovacs 1997; Houser 2025; Reichmuth et al 2013; Thewissen and Nummela 2008 [14–16,41] |
| *Spilogale putorius* | In-air only | Above water | Terrestrial | Nowak 2003 [13] |
| *Tasmacetus shepherdi* | Underwater only | NA | NA | Nowak 2003; Thewissen and Nummela 2008 [13,14] |
| *Taxidea taxus* | In-air only | Above water | Terrestrial | Nowak 2003 [13] |
| *Tremarctos ornatus* | In-air only | Above water | Terrestrial | Nowak 2003 [13] |
| *Tursiops truncatus* | Underwater only | NA | NA | Nowak 2003; Thewissen and Nummela 2008 [13,14] |
| *Ursus americanus* | In-air only | Above water | Terrestrial | Nowak 2003 [13] |
| *Ursus arctos* | In-air only | Above water | Terrestrial | Nowak 2003 [13] |
| *Ursus maritimus* | In-air only | Shallow | Terrestrial | Nowak 2003; Thewissen and Nummela 2008; Stirling and van Meurs 2015; Lone et al 2018 [13,14,42,43] |
| *Ursus thibetanus* | In-air only | Above water | Terrestrial | Nowak 2003 [13] |
| *Vormela peregusna* | In-air only | Above water | Terrestrial | Nowak 2003 [13] |
| *Vulpes lagopus* | In-air only | Above water | Terrestrial | Nowak 2003 [13] |
| *Vulpes vulpes* | In-air only | Above water | Terrestrial | Nowak 2003 [13] |
| *Zalophus californianus* | Amphibious | Mesopelagic | Aquatic | Schreer and Kovacs 1997; Houser 2025; Reichmuth et al 2013; Thewissen and Nummela 2008 [14–17] |
| *Zalophus wollebaeki* | Amphibious | Mesopelagic | Aquatic | Villegas-Amtmann et al 2008; Houser 2025; Reichmuth et al 2013; Thewissen and Nummela 2008 [14–16,44] |

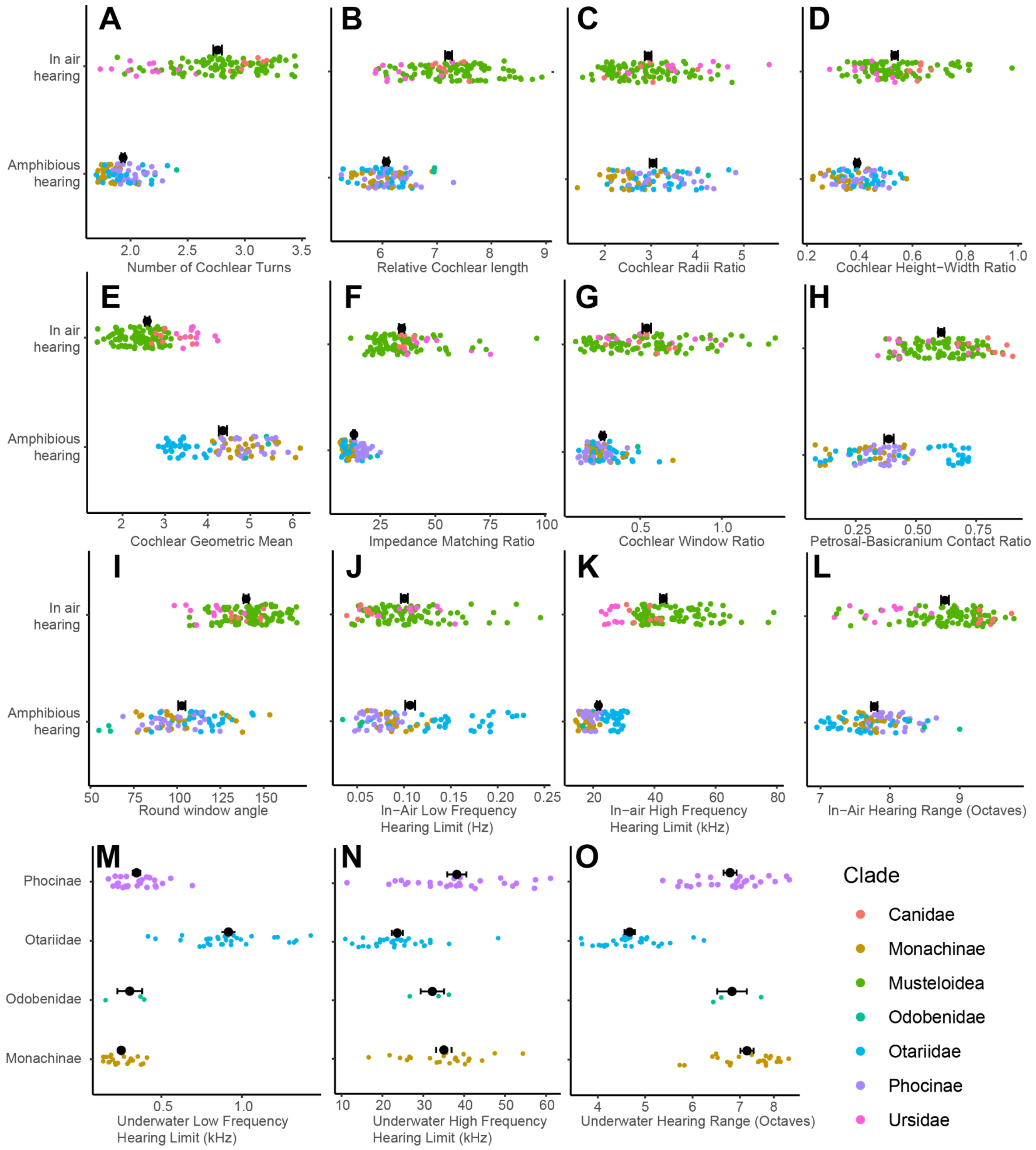


**Figure S2. Morphometric and auditory data of extant caniform species, related to Table 1 and Methods.** (A) Number of cochlear turns, (B) scaled cochlear length , (C) cochlear radii ratio, (D) cochlear height-width ratio, (E) cochlear geometric mean, (F) impedance matching ratio (tympanic membrane area to oval window area), (G) cochlear window ratio (oval window area to round window area, (H) petrosal contact ratio (contact between basicranium and petrosal perimeter), (I) angle between round window and main axis of petrosal, (J) in-air low frequency hearing limit (kHz), (K) in-air high frequency hearing limit (kHz), (L) in-air hearing range (octaves), (M) underwater low frequency hearing limit (kHz), (N) underwater high frequency hearing limit (kHz), (O) underwater hearing range (octaves). Black points and error bars represent the mean and standard error range.

**Table S3. Phylogenetic signal test from Methods and Results.** $Log_{10}$ morphometric and bioacoustic traits.

| Trait | Blomberg's K | Pagel's λ |
|---|---|---|
| $Log_{10}$ Cochlear Turns | 0.79 | 0.92 |
| $Log_{10}$ Relative Cochlear Length (mm) | 0.33 | 0.8 |
| $Log_{10}$ Cochlear Radii Ratio | 0.16 | 0.41 |
| $Log_{10}$ Cochlear Height-Width Ratio | 0.21 | 0.86 |
| $Log_{10}$ Impedance Matching Ratio | 0.68 | 0.9 |
| $Log_{10}$ Cochlear Window Ratio | 0.11 | 0.50 |
| $Log_{10}$ Petrosal-Basicranium Contact Ratio | 0.1 | 0.59 |
| $Log_{10}$ Round Window Angle (°) | 0.37 | 0.91 |
| $Log_{10}$ Low Frequency Hearing Limit (kHz) | 0.2 | 0.85 |
| $Log_{10}$ High Frequency Hearing Limit (kHz) | 1.44 | 0.97 |
| $Log_{10}$ Octaves | 0.37 | 0.88 |

**Table S4. Estimates of hearing mode for each extinct pinniped species used in Flexible Discriminate Analysis in Figure 2.** Values represent posterior probabilities of each hearing mode. M+I: Middle and Inner ear model (BGVE: 100%; df: 9; TME: 0%); MO: Middle ear only model (BGVE: 100%; df: 5; TME: 1.04%); IO: Inner ear only model (BGVE: 100%; df: 5; TME: 8.8%). *Alternate middle ear only model which removes the petrosal contact ratio, not preserved in *Enaliarctos mitchelli* specimen (BGVE: 100%; df: 4; TME: 1.04%). N/A indicates a model non-applicable for taxa lacking the full preserved morphology. BGVE: between-group variance explained; DF: degrees of freedom; TME: training misclassification error.

| Species | Middle and Inner ear model | | Middle ear only model | | Inner ear only model | |
|---|---|---|---|---|---|---|
| | In-air only | Amphibious | In-air only | Amphibious | In-air only | Amphibious |
| *Acrophoca longirostris†* | <0.01% | >99.99% | <0.01% | >99.99% | 1.56% | 98.44% |
| *Allodesmus kernensis†* | <0.01% | >99.99% | <0.01% | >99.99% | 1.49% | 98.51% |
| *Callorhinus* sp† | N/A | N/A | N/A | N/A | 1.83% | 98.17% |
| *Desmatophoca brachycephala†* | N/A | N/A | N/A | N/A | 1.25% | 98.75% |
| *Devinophoca claytoni†* | <0.01% | >99.99% | <0.01% | >99.99% | 0.83% | 99.17% |
| *Enaliarctos mealsi†* | N/A | N/A | N/A | N/A | 23.16% | 76.84% |
| *Enaliarctos mitchelli†* | N/A | N/A | <0.01%* | >99.99%* | N/A | N/A |
| *Eomonachus belegaerensis†* | <0.01% | >99.99% | <0.01% | >99.99% | 4.62% | 95.38% |
| *Hadrokirus martini†* | N/A | N/A | <0.01% | >99.99% | N/A | N/A |
| *Homiphoca capensis†* | <0.01% | >99.99% | <0.01% | >99.99% | 0.65% | 99.35% |
| *"Leptophoca lenis"* CMM V 2021*†* | <0.01% | >99.99% | <0.01% | >99.99% | 1.18% | 98.82% |
| *Neomonachus tropicalis†* | <0.01% | >99.99% | <0.01% | >99.99% | 2.27% | 97.73% |
| *Neotherium mirum†* | <0.01% | >99.99% | <0.01% | >99.99% | 13.10% | 86.90% |
| *Pinnarctidion iverseni†* | <0.01% | >99.99% | <0.01% | >99.99% | 9.07% | 90.93% |
| *Pinnarctidion* sp.*†* | N/A | N/A | N/A | N/A | 20.01% | 79.99% |
| *Piscophoca pacifica†* | <0.01% | >99.99% | <0.01% | >99.99% | 3.75% | 96.25% |
| *Pithanotaria starri†* | <0.01% | >99.99% | 0.55% | 99.45% | 0.17% | 99.83% |
| *Pontolis barroni†* | N/A | N/A | N/A | N/A | 23.43% | 76.57% |
| *Potamotherium vallentoni†* | 0.39% | 99.61% | 99.86% | 0.14% | 38.21%% | 61.79%% |
| *Prototaria planicephala†* | <0.01% | >99.99% | 0.17% | 99.83% | 2.47% | 97.53% |
| *Pteronarctos goedertae†* | N/A | N/A | 1.42% | 98.58% | N/A | N/A |
| *Puijila darwini†* | 14.36% | 85.64% | 0.89% | 99.11% | 28.36% | 71.64% |
| *Sarcodectes magnus†* | <0.01% | >99.99% | <0.01% | >99.99% | 35.58% | 64.42% |
| *Zalophus japonicus†* | <0.01% | >99.99% | <0.01% | >99.99% | 0.80% | 99.20% |

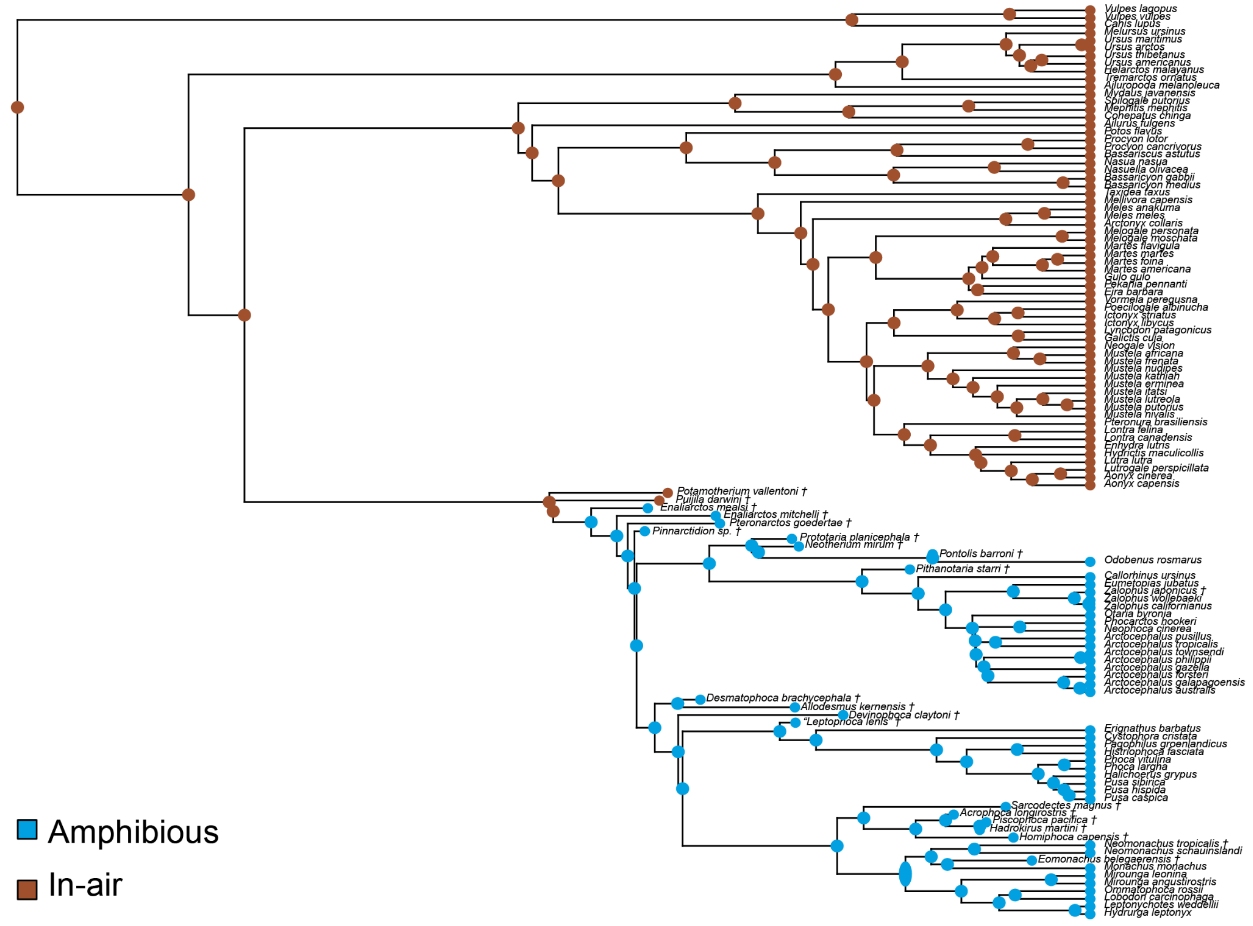


**Figure S3. Ancestral state estimation from Figure 2.** Full results for equal-rates analysis.

**Table S5. Results of comparative trait modelling in BayesTraits related to Figure 3.** We ran three independent chains of each model for each trait and compared model fit using marginal likelihoods (indicated below). In the variable rates model, traits are allowed to evolve via individual branch-specific rates which depart from a background rate of trait evolution. The variable rates model is preferred for all traits, with the best model fit highlighted in grey.

| | Brownian motion | | | Variable rates | | | Early burst | | | Ornstein–Uhlenbeck | | | |
|---|---|---|---|---|---|---|---|---|---|---|---|---|---|
| **Trait** | **Run 01** | **Run 02** | **Run 03** | **Run 01** | **Run 02** | **Run 03** | **Run 03** | **Run 02** | **Run 01** | **Run 01** | **Run 02** | **Run 03** | **Best Fit** |
| Cochlear Turns | 85.49 | 85.44 | 85.54 | 122.09 | 122.24 | 122.08 | 86.34 | 85.79 | 86.05 | 87.09 | 87.31 | 87.45 | VarRates |
| Relative Cochlear Length | 99.98 | 99.90 | 100.24 | 120.07 | 120.26 | 120.14 | 99.48 | 99.99 | 99.51 | 103.55 | 103.73 | 103.73 | VarRates |
| Cochlear Radii Ratio | -20.93 | -20.92 | -20.94 | -7.29 | -7.31 | -7.39 | -19.87 | -19.78 | -19.81 | -22.35 | -22.14 | -22.00 | VarRates |
| Cochlear Height-Width Ratio | -12.48 | -12.56 | -12.35 | 17.20 | 17.34 | 17.11 | -12.74 | -12.73 | -12.83 | -8.31 | -8.37 | -8.27 | VarRates |
| Impedance Matching Ratio | -73.17 | -73.23 | -73.19 | -47.47 | -47.46 | -47.94 | -67.32 | -67.25 | -67.30 | -77.63 | -77.82 | -77.73 | VarRates |
| Cochlear Window Ratio | -126.49 | -126.53 | -126.45 | -93.47 | -93.60 | -93.45 | -127.27 | -126.82 | -127.22 | -121.32 | -121.22 | -121.20 | VarRates |
| Petrosal-Basicranium Contact Ratio | -113.42 | -113.38 | -113.45 | -35.55 | -35.40 | -35.35 | -113.82 | -115.02 | -114.61 | -96.57 | -96.65 | -96.70 | VarRates |
| Round Window Angle | 17.49 | 17.36 | 17.39 | 60.05 | 60.11 | 59.96 | 19.00 | 19.08 | 19.14 | 15.99 | 16.05 | 15.89 | VarRates |
| In-air Low Frequency Hearing Limit (kHz) | -59.93 | -59.76 | -59.76 | -36.51 | -36.54 | -36.57 | -59.78 | -59.69 | -59.75 | -56.50 | -56.50 | -56.38 | VarRates |
| In-air High Frequency Hearing Limit (kHz) | 40.67 | 40.80 | 40.76 | 53.77 | 54.03 | 54.52 | 43.61 | 43.50 | 43.65 | 37.42 | 37.53 | 37.48 | VarRates |
| In-air Hearing Range (Octaves) | 145.54 | 145.69 | 145.54 | 188.61 | 188.62 | 188.54 | 145.82 | 145.63 | 145.50 | 150.69 | 150.71 | 150.70 | VarRates |
| Underwater Low Frequency Hearing Limit (kHz) | -38.22 | -38.24 | -38.19 | -27.60 | -27.54 | -27.56 | -38.38 | -38.34 | -38.27 | -40.45 | -40.37 | -40.26 | VarRates |
| Underwater High Frequency Hearing Limit (kHz) | -55.98 | -55.89 | -55.86 | -45.70 | -45.63 | -45.66 | -52.21 | -52.12 | -52.25 | -59.54 | -59.54 | -59.61 | VarRates |
| Underwater Hearing Range (Octaves) | -16.04 | -16.07 | -16.10 | -1.22 | -1.30 | -1.23 | -10.93 | -10.87 | -10.97 | -19.98 | -19.91 | -19.94 | VarRates |

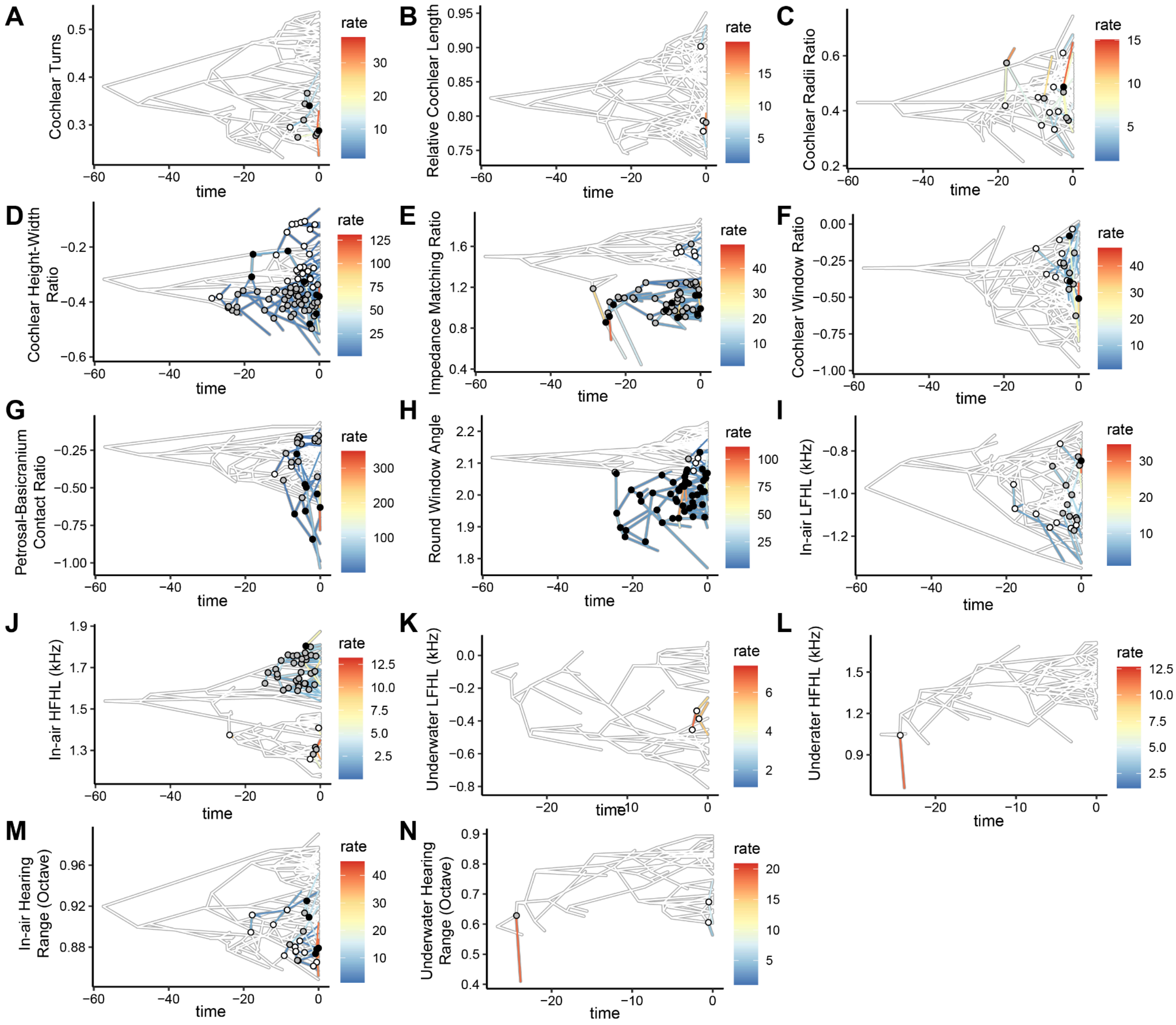


**Figure S4. Variable rate shifts of morphometric and auditory traits from Figure 3 ($log_{10}$ transformed).** (A) Cochlear turns, (B) relative cochlear length, (C) cochlear radii ratio, (D) cochlear height-width ratio, (E) impedance matching ratio, (F) cochlear window ratio, (G) petrosal-basicranium contact ratio, (H) round window angle, (I) in-air Low Frequency Hearing Limit (LFHL), (J) in-air High Frequency Hearing Limit (HFHL), (K) underwater LFHL, (L) underwater HFHL, (M) in-air hearing range (octaves), (N) underwater hearing range (octaves). Circle colours: White = putative shifts; grey = likely shifts; black = positive selection [45]. Evolutionary rates are presented as rates relative to this background rate and vary considerably across traits. For example, some traits are comparatively conservative such as the in-air High Frequency Hearing Limit (in kHz) which shows limited departures from the background rate but includes both rate increases (up to ~10x) and decreases (as little as 0.25x). In comparison, other traits show rapid and dramatic changes in rates (up to 200x) such as in the evolution of the petrosal-basicranium contact ratio, particularly in the otariid genus *Zalophus*. Results from tests for variable rate shifts can be found in the Figshare repository.

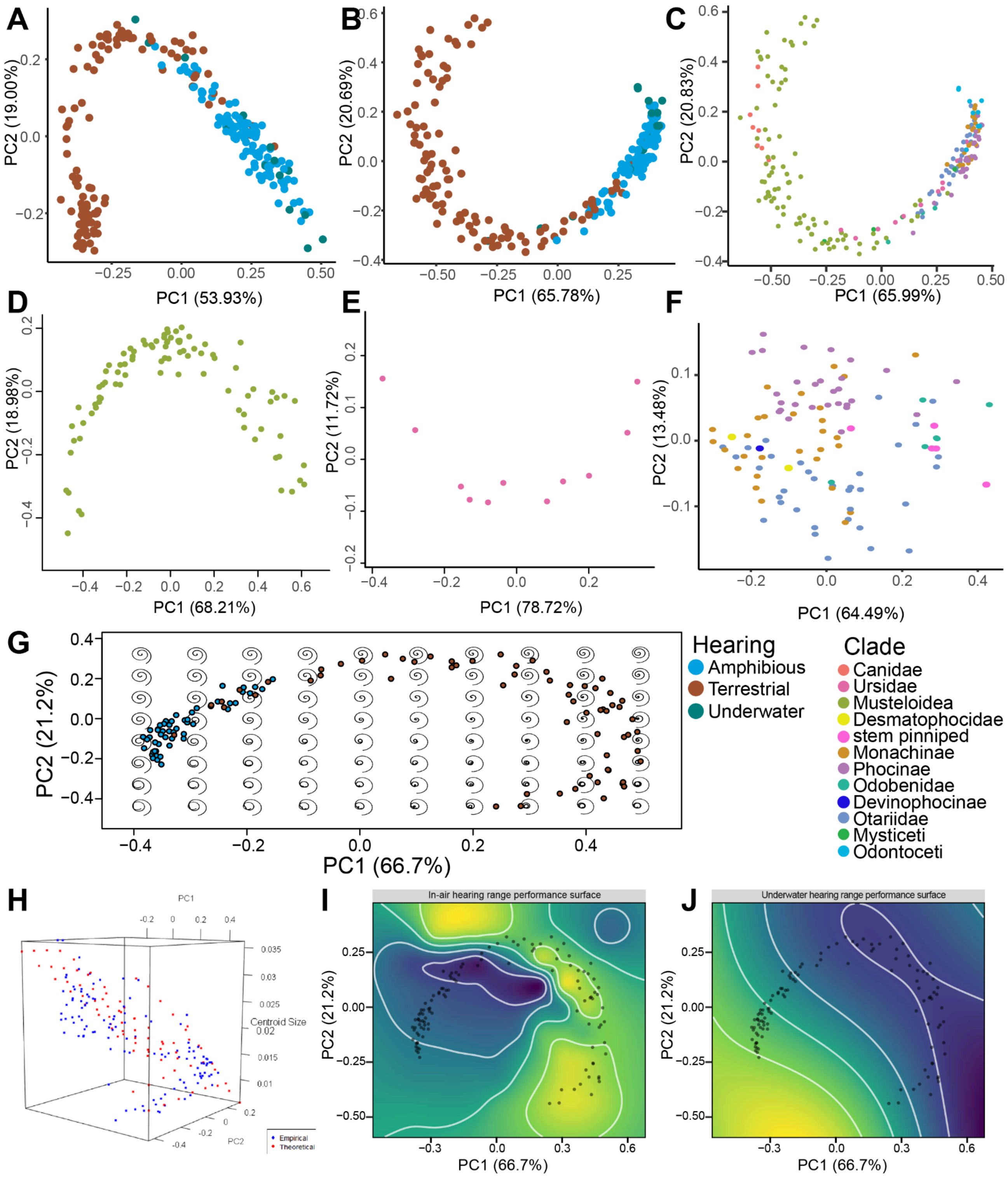


**Figure S5. 3D geometric morphometric analyses relating to Figure 5**. Principal components analysis (PC1 and PC2 morphospaces) of generalised Procrustes analysis (GPA) of landmark data of the inner ears of Caniformia (and cetaceans) specimens. (A) Caniformia and Cetacea with sliding semilandmarks. (B) Caniformia and Cetacea with fixed landmarks. (C) Extant Caniformia with fixed landmarks. (D) Musteloidea with fixed landmarks. (E) Ursidae with fixed landmarks (F) Pinnipedia with fixed landmarks. (G) Projection of 80 theoretical cochlea shapes onto the morphospace. (H) projection of predicted centroid size onto theoretical cochlea shapes, used for calculating hearing range. Performance surfaces of theoretical (I) in-air and (J) underwater hearing ranges. See code in Figshare for details on GPA and PCA.

#### Method S1. Micro-CT scanning

Most new specimens were scanned on a Nikon Metrology HMX ST 225 Micro-CT scanner at the Imaging and Analysis Center, Natural History Museum London, and the Molecular Imaging Center, University of Southern California. Specimen resolution was assessed on a case-by-case basis, (voxel size between 6-150 microns) to ensure the entire auditory region was in view. See Morphosource Project (ID 000656781: SEAL: The Evolution of Auditory Adaptations for Aquatic Life in Pinnipeds) for scan settings. Scans were processed in Dragonfly (Version 2021.1) or Avizo (version 2021.2). The middle earbone was segmented out, and for the inner ear the negative space of the cochlea was segmented out. Fossil taxa mostly required manual segmentation, with either/both the middle and inner ear segmented depending on scan quality. Meshes were smoothed in Avizo or Dragonfly before exporting as .ply files. Once created, excess material was removed in Meshlab (Version 2022.02) and meshes were downsampled by 50% (if needed).

#### Method S2. Data measurements

All measurements were taken in millimetres. For a subset of specimens scanned at the Natural History Museum London, (i) anterior-posterior skull length (anterior premaxilla to ventral base of occipital condyle), (ii) dorsal-ventral skull height (anterior-posterior height of occipital bone), (iii) maximum lateral skull width (lateral most width of skull at zygomatic arches), (iv) tympanic bulla anterior-posterior length, and (v) tympanic bulla medial-lateral width, were taken to assess allometry. To test if earbone size scales with skull size, we regressed the geometric mean of skull measurements (length, width, height), against the geometric mean of tympanic bulla measurements (length, width). An $R^2$ >80 meant that the geometric mean of the tympanic bulla could be used to scale middle ear measurements (see Figure 4a). Linear measurements chosen for the middle ear were informed based on observations first noted in Repenning [46], and the inner ear based on West [47], Manoussaki [48], and Taszus et al. [1].

***Impedance Matching Ratio and Window Ratio*****:** Mammals conduct sound to the inner ear via the tympanic membrane, middle ear ossicles, and oval window. Middle ears amplify airborne sound signals so that they overcome the impedance mismatch between the air sound travels in and the fluid of the cochlea: this is achieved through the lever arm of the middle ear ossicles, and the enlarged size of the tympanic membrane relative to the oval window (the impedance matching ratio).

Mammalian auditory anatomy relies on air, both in the ear canal and the middle ear cavity to function. However, underwater the mammalian ear does not function, due to the impedance mismatch between the air in the ear and the water sound travels in. Pinnipeds, which have mammalian ears but can hear in both environments, are therefore puzzling. One theory put forward by Repenning [46] for how pinnipeds overcome this limitation is by receiving sounds via the cavernous tissue in their ear canal and middle ears. This cavernous tissue flushes with blood to limit the space air occupies during diving and equalise ear pressure. By doing this, pinnipeds fill their auditory pathway with a fluid filled substance (the blood in the tissue) closer to the impedance of the water sound travels through. If this is the case, a middle ear amplifying underwater noises (which are louder and more powerful than in air) risks damaging the inner ear. Therefore, it would be expected that pinnipeds have a lower tympanic membrane to oval window size ratio.

Based on this, as well as observations by Repenning [46] and Taszus et al. [1], we measured the impedance matching ratio of Caniformia to test for this (Figure S1). Both studies also report that some pinnipeds possess a large round window relative to their oval window (cochlear window ratio), which may also be important for underwater hearing. To measure these ratios we used the following protocol: meshes of the full earbones were imported into meshlab, and the view was cropped so the middle ear cavity was visible. Maximum length and perpendicular width were taken using the measure tool for the tympanic sulcus, oval window, and round window. The ovoid area of each of these structures was then calculated using the following equation:

$$Ovoid\ area = \pi \times maximum\ length \times perpendicular\ width$$

Then, the Impedance Matching Ratio (IMR) was calculated by dividing the tympanic membrane area by the oval window area. The Window Ratio (WR) was calculated by dividing the oval window area by the round window area.

***Petrosal bone contact ratio*:** Fully aquatic cetaceans, which have excellent underwater hearing, have limited the contact of their earbones with the surrounding basicrania. This is thought to help limit noise being detected in the cochlea via bone conduction from the basicranium. Pinnipeds, particularly phocids, have been observed to have limited contact between the petrosal and the basicranium [46,49], and even the bony tentorium (which normally contacts the petrosal dorsally) has been retracted from the ear.

We therefore quantified the bone contact ratio between the petrosal and the basicranium (Figure S1) via the following protocol: Full earbone meshes were imported into the program MorphoDig, and aligned so that the dorsal view of the petrosal was visible. Then, landmarks were placed along the perimeter of the petrosal from the medial contact with the basicranium at the level of the floccular fossa, to the lateral contact with the basicranium at the level of the floccular fossa. Using these landmarks, a curve of 50 equidistant landmarks was created, and its length measured for the petrosal perimeter. Then, all landmarks that were not placed where the petrosal contacted the basicranium were deleted, and any uninterrupted lengths of the curve had their lengths calculated, and added to give the bone contact length. Then, the petrosal perimeter was divided by the bone contact length to give the bone contact ratio.

***Round Window Angle*:** The primary function of the round window is to dissipate the sound energy, which travelled from the movement of the oval window through the cochlea, into an air filled chamber in the tympanic bulla. Normally, the round window is orientated to face this chamber. But in Phocidae, the round window is not only enlarged, but also faces away from the tympanic cavity completely; in some species, it abuts the posterior tympanic bulla wall; in others, the tympanic bulla opens up into a foramen (the external cochlear foramen), so that the round window can be seen from outside of the skull [46,50]. While Taszus et al. [1] discuss an external cochlear foramen in detail, the morphology they identify is not the external cochlear foramen, and it is instead the cochlear aqueduct. This reorientation of the round window is possibly an adaptation moving the window away from the cavernous tissue in the middle ear [46]. This is important, because if the cavernous tissue conducts sound, then any sound exiting the cochlea via the round window would travel through the cavernous tissue and potentially restimulate the oval window again. We therefore measured the angle of the round window against the petrosal (Figure S1) using the following protocol.

Full earbone meshes were cropped in Meshlab so that the tympanic bullae was removed, and the round window was completely visible. These meshes were imported into Rhino 3D, and the position was rotated so that the medio-lateral and anterior-posterior axis of the petrosal was aligned with the z-axis. Then, a 2D plane was established roughly at the flat axis (medio-lateral and anterior-posterior) of the petrosal. It was calculated using three landmarks: one landmark dorsal to the canal for *M. tensoris veli Palatini*, between the petrosal and the basicranium; one landmark at the anterior apex of the petrosal; and one landmark at the anterior-dorsal opening of the cochlear aqueduct. A second 2D plane was then established using the three more landmarks: the medial, dorsal, and lateral most points of the round window fossula (opening). Then, a planar angle was taken between these two planes.

***Cochlear geometric means and Height-width ratio***: To scale inner ear measurements, and to test 3D cochlear shape for allometry, geometric means were calculated from cochlear measurements of the 3D landmarks of the "dorsal" midline of the cochlea. Three measurements were using the "measure" tool in Rhino 3D v.8 for the geometric means: cochlear width, defined as the maximum width of the cochlea at the outer turn, taken parallel to the round window; cochlear depth, defined as the maximum depth of the cochlea taken perpendicular to the width; and cochlear height, defined as the maximum "dorso-ventral" height of the cochlea. Taszus et al. [1] observed that a "tower" morphology for the cochlea was present in some terrestrial Caniformia that is absent in pinnipeds. A cochlear height-width ratio was therefore used to test this morphology (Figure S1). The cochlear height measurement was divided by the cochlear width measurement to provide the cochlear height-width ratio (basal ratio).

***Cochlear Length***: The length of the basilar membrane has been observed to be important for determining the sounds tetrapods can hear [12,47,51], and we therefore included cochlear length (as a proxy) in our analyses. From the 100 equidistant semi-sliding landmark curve from the prior step, the length of this 3D curve was calculated using the "measure curve" tool in MorphoDig. The 3D length of theoretical cochlear shapes was calculated using the "length" tool in Rhino 3D V.8. Cochlear length was scaled to cochlea size by dividing it by the geometric mean of the cochlea (width, depth, height) defined above.

***Cochlear turns***: Therian mammals have coiled cochlea, an adaptation that helps maximise cochlear length while saving space in the petrosal [12,47]. The number of turns of the cochlea is potentially related to hearing frequency, with a small number of turns related to maximum hearing frequency, and a larger number of turns related to minimum hearing frequency [47,48].

Landmarks were imported into Rhino 8 to measure the number of turns (Figure S1). First, .pts landmark files were converted to .txt files using the command prompt. Then, landmarks were imported into Rhino 8, with the grid scale altered to match the landmark scale (e.g. microns). A z-plane was established on the X-Y axis. The rotate function was used to move the landmarks so that the beginning of the first turn is horizontal to the x-axis. Next, a point was placed at the apex of the cochlea (end landmark). Then, lines were established directly horizontal of this point (in the top view), and in the orientation the apex is pointing towards. Using the angle dimension function, the angle was measured between these two lines, from the bottom of the horizontal line to the apex line. Dividing the angle by 360, and adding the number of prior turns results in the turn number. For some theoretical shapes exported after the 3D geometric morphometric analysis, the middle turn folds in on itself. This is a product of how new turns are generated during Procrustes superimposition. We treated these turns as partial turns, calculating their perimeter along the 2D flattened curve, and dividing it by the perimeter (excluding the infolding) of the turn it was folding within, which was then added to the turn number.

***Radii Ratio***: The ratio between the radius of the turn at the base of the cochlea (R base) and the radius of the turn at the end of the cochlea (R apex) is thought to be related to minimum hearing frequency limit [48]. In addition, it is needed for underwater hearing frequency calculations (see Methods below).

Landmarks were copied to a new layer called Place on Plane, with the original landmarks hidden. These points were then flattened onto a z-plane using the function ProjectToCPlane, retaining their x and y coordinate positions. The CurveThroughPt function was used to calculate a curve between the landmark points in a layer called Draw curve. Previous layers were hidden so only Draw Curve is visible. A point is placed at the start at the curve and the end of the curve. Then a line perpendicular to both these points is drawn that ends parallel to a quarter turn. Then, a line perpendicular to the end of those lines (so it is 90 degrees from the prior line) is drawn to intersect with the curve, and a new point is established at this intersection. The split function is used on the curve with the points as cutting objects. This will provide the first quarter turn and last quarter turn, which can be copied into new layers. The divide function is then used to split the first ¼ turn and last ¼ turn of the cochlea into four sections. For each ¼ turn segment, a line was drawn between points 1-3, and 3-5. These lines were divided in two, then a line was drawn perpendicular from the midpoint of these two lines. A line was drawn between the intercept of those two lines, and point 3 on the curve. Measuring the length of these two lines using the length function provides both Radii Base and Radii Apex measurements. Then the Radii Base was divided by the Radii Apex to get the Radii Ratio (Figure S1).

***In-Air Hearing Frequencies***: The underwater hearing abilities of pinnipeds are thought to come at the cost of a limited range of in-air hearing when compared to their terrestrial relatives [16]. In order to provide an estimate of relative in-air hearing frequency limits for each specimen, the following equations from West [47] were used.

$$Log10\ LFHL\ in\ Air\ (60dB\ re\ 20\ \mu Pa\ [SPL]) = 1.76 - 1.66 \times Log10(Cochlear\ length \times spiral\ turns)$$

$$Log10\ HFHL\ in\ Air\ (60dB\ re\ 20\ \mu Pa\ [SPL]) = 2.42 - 0.99 \times Log10(Cochlear\ length \div spiral\ turns)$$

Where LFHL is Low Frequency Hearing Limit and HFHL is High Frequency Hearing Limit (in kHz), dB is decibels, μPa is micro pascals, and SPL is sound pressure level.

While the Low Frequency Hearing Limit equation from Manoussaki et al. [48] is usually preferred over the West [47] equation, we opted for the latter as it enables the Low and High frequency hearing limit estimates to be comparable due to using the same data and measurements in their equations. This is necessary, as no equation can currently estimate hearing frequency limits accurately, and it is best to provide estimates that give a relative estimation between individuals in the sample.

***Underwater Hearing Frequencies***: There are currently no methods to estimate underwater hearing range in mammals. We developed two equations to estimate the region of best hearing underwater using the pinniped data contained in Table 8.1 of Houser [15], and species mean morphometric data from this study. The below equations are reminiscent of those from West [47], are designed to provide a means of calculating relative hearing range for comparison, and are by no means accurate predictions.

$$Log10\ LFHL\ Underwater\ (kHz) = -1.3798 \times Log10\left(\frac{(Cochlear\ length \times RApex)}{spiral\ turns}\right) + 1.2294$$

Average Percent Prediction Error = 27.51%
Percent Standard Error of the Estimate = 48.83%

$$HFHL\ Underwater\ (kHz) = 11.931 \times \left(\frac{scaled\ cochlear\ length \times \left(\frac{RBase - RApex}{spiral\ turns}\right)}{spiral\ turns}\right) - 12.688$$

Average Percent Prediction Error = 16.14%
Percent Standard Error of the Estimate = 45.16%

While these equations work for pinnipeds and cetaceans, they do not perform well for most in-air only hearing species and some theoretical cochlea. This is because they either produce HFHL lower than LFHL, or negative values. In these cases, the underwater hearing values are instead substituted with 0, and it is assumed these cochlea are deaf to underwater sounds. Once hearing frequency limits were estimated, hearing range was calculated in octaves via a transposed version of the standard equation:

$$Hearing\ range\ (Octaves) = Log10(HFHL \div LFHL) \div Log10(2)$$

##### Method S3. Evolutionary Modelling

To explore the evolution of each individual morphometric and auditory trait, we fit and compared a set of evolutionary models using BayesTraits. These models covered unbiased random walks (Brownian Motion), single optimum Ornstein-Uhlenbeck (OU), accelerating or decelerating rates through time (EB), and a variable rates model which allows branch-specific rates. The species averaged data and the phylogenetic tree from prior analyses were used. The variable rates model allows each branch to have its own Brownian motion rate, and has been shown to outperform other models when exploring the role of evolutionary bursts toward novel morphologies [52].

We fit three independent chains of each model to assess convergence, and estimated marginal likelihoods using a stepping-stone sampler (500 stones each for 5,000 generations). BayesTraits V4 implements a reversible-jump MCMC sampler, which we specified to run for 100 million generations with 10 million generations of burnin. We summarized the output to check for convergence with custom scripts and verified that all parameters had reached effective sample sizes (ESS) > 200. To verify model fit and consistency we compared results across chains. Individual model outputs were summarized using the Variable Rates Post Processor software.

To track trait values at internal nodes we logged values from the MCMC chain using the *AddTag* and *AddMRCA* features in BayesTraits. We summarized posterior estimates of evolutionary rates (mean, median) along individual branches. The relative evolutionary rates of the best supported model can then be used to explore shifts in the evolution of each trait. We visualized branch-specific evolutionary rates with custom scripts (available in the supplement). This is useful when exploring how selective pressures (such as the transition from land to water) resulted in novel phenotypes (amphibious hearing mode).

##### Method S4. 3D geometric morphometric analyses

Cochlear meshes were imported into Morphodig (Version 1.6.9), and aligned with the round window facing forward. Landmarks were placed onto the “dorsal” surface of the cochlea (defined as the orientation at which the apex of the cochlea is in full view, Figure S1) at the midline of the duct, starting from the apex of the cochlea and ending at the base of the cochlea, where the cochlear duct intersects with the ampullae. These landmarks were then exported, and reimported as handles, and the landmarks were rotated by 20% semi-automatically to optimise the curve to the mesh surface. Landmarks were resampled as 100 equidistant landmarks. For analyses where landmarks were slid, a sliding matrix was created, with landmark 1 and 100 as fixed landmarks, and landmarks 2-99 as sliding landmarks (but see below about why this may not be appropriate).

To assess whether the 3D shape of the cochlea varies by hearing mode (amphibious, terrestrial only, underwater only) we performed a 3D geometric morphometric (GMM) analysis of all specimens for the 100 equidistant landmarks exported from MorphoDig. We ran one analysis treating the landmarks as semi-sliding along the curve, and one with them as all fixed landmarks (see below). The Generalised Procrustes Analysis of landmarks was done using the gpagen function in the R packaged geomorph. The principal component analysis of the 3D coordinates after Procrustes superimposition was done with the gm.prcomp function in geomorph. We also created morphospaces for specific clades (musteloids only, ursids only, and pinnipeds only) to explore how cochlear shape varied within clades. We also repeated analyses with mean shape data (and accompanying mean metadata) for all species. We generated morphospaces which were plotted using the R package ggplot2.

Normally for a landmark curve, landmarks are slid using a sliding matrix prior to analysis [53,54]. For our analysis, this would involve having landmarks 1 and 100 fixed, and sliding 2-99. This is usually done to minimise bending energy along the curve, and is a common practice among 3D GM analyses which quantify the cochlea as a single curve [1,55–57]. However, this effectively collapses the majority of the landmarks along the first turn, which is the largest turn on the outside of the spiral. As a result, any analysis done after sliding would be on a shape that has minimal landmarks representing the inner turns, limiting downstream interpretations. Some studies [55,56] modify the sliding settings in an attempt to address this issue, but this only partially resolves this issue after Procrustes superimposition. To address this, we first did analyses without sliding, as once landmarks are resampled to be equidistant, homology along the length of the cochlea is already established (as a cochlea with more turns would be the result of elongation of the cochlear duct). We then repeated analyses with sliding. This produced similar results for the morphospace and statistical tests (see Figshare repository for statistical tests).

We also recommend caution in the interpretation of 3D GM morphospaces which analyse the cochlea as a single curve. After Procrustes superimposition, the principal component analysis of a single curve effectively results in a highly restricted morphospace occupation. This morphospace is seen in several analyses [1,55–57], as well as our own (below). This is the result of the movement of the landmarks between each specimen from the mean shape and other specimens being strictly linear between PC axes. As a result, morphospace occupation between PC axes is restricted, and shapes should be checked for impossible configurations (e.g. an anti-clockwise cochlea) after Procrustes superimposition. While these morphospaces provide useful shape differences related to the coiling of the cochlea, absolute cochlear shape differences (e.g. relative height, relative width) will be reduced. We mention this to stress the importance of checking landmark configurations after Procrustes superimposition.

Finally, we used theoretical morphospace methods on the species mean morphospace to generate performance surfaces and adaptive landscapes for the auditory variables in-air hearing range in octaves and underwater hearing range in octaves using the R packages morphospace and Morphoscape. This enabled us to assess whether the evolution of underwater hearing in pinnipeds limited in-air hearing (as observed in Reichmuth et al 2013 [16]) and cochlear shape. A theoretical morphospace was created using the mspace function by sampling 80 theoretical shapes (10 columns, 8 rows) from PC1 and PC2. These theoretical landmark coordinates were then exported, and imported into Rhino3D version 8 to take inner ear measurements using the protocol used for the actual cochlear landmarks. We used the empirical cochlear shapes to regress centroid size against PC1 x PC2 x cochlear turns, and the resulting linear model was used with the predict function in R to estimate centroid size for the 80 sampled theoretical shapes. We then multiplied the estimated centroid size of each theoretical shape against the morphometric data measured in Rhino3D, and used the resulting scaled measurements to estimate theoretical auditory data using the previous equations for in-air and underwater hearing frequency limits.

The auditory data for the theoretical shapes was then imported back into R. The krige_surf function was used to generate performance surfaces. and the calcWprimeBy function was used with generated performance surfaces to calculate adaptive landscapes by hearing mode (Terrestrial vs Amphibious). We did this for theoretical hearing ranges (in-air octaves, underwater octaves), empirical hearing ranges, and theoretical + empirical hearing ranges.
